# SureCLIP and DrCLIP: Genetic manipulation-free, sub-hour protein interaction mapping in cells and zebrafish

**DOI:** 10.64898/2025.12.04.692439

**Authors:** Brady T. Cline, Chad D. Torrice, Weihua B. Fan, Sumaya A. Addish, Mark Hazelbaker, Heejun Lee, James E. Bear, Ukrae H. Cho

**Author notes:** Correspondence should be addressed to Ukrae H. Cho.

## Abstract

Proximity labeling has made interactome profiling broadly accessible, but it requires transgenic cell lines and animals. In primary or sensitive cells, deriving lines risks clonal drift and overexpression artifacts. In zebrafish, transgenesis imposes at least six months of lead time, and overnight labeling cannot resolve events on the timescale of embryogenesis. Here, we present SureCLIP and DrCLIP, protein interaction mapping methods that achieve sub-hour temporal resolution in cultured cells and zebrafish, respectively, without genetic manipulation. Central to these methods are the reversible crosslinkers DSP and DTBP. SureCLIP analysis of Hic-5 and lamin B1 compared favorably with published proximity labeling datasets in coverage and enrichment and identified new interactors. DrCLIP enabled stage-resolved endogenous interactome profiling in wild-type zebrafish and revealed that kindlin-2 is required for ECM assembly at somite boundaries and for myotendinous junction formation. Together, SureCLIP and DrCLIP circumvent the logistical, biological, and temporal limitations of transgene-based protein interaction mapping.

## INTRODUCTION

Interactome mapping approaches fall into two main categories: (1) co-immunoprecipitation coupled with mass spectrometry (co-IP/MS)^1,2^ and (2) proximity labeling (e.g., BioID, TurboID, and APEX)^3–5^. The latter has become increasingly popular, partly because it requires far less optimization. In co-IP/MS, one must identify lysis and wash conditions that are strong enough to extract proteins and remove non-specific binders, yet gentle enough to preserve protein-protein interactions (PPIs) throughout. Proximity labeling circumvents the dilemma by covalently tagging interactors prior to lysis.

However, this comes at a cost. Bait-enzyme fusions must be genetically introduced, which lengthens experimental timelines, compromises biological fidelity, and limits temporal resolution. First, generating transgenic cell lines or model organisms adds weeks to months to the workflow. For zebrafish, following F0 injection, obtaining sexually mature, non-mosaic F1 carriers takes at least half a year. Second, this approach risks genetic and phenotypic drift, a well-documented concern in primary cultures^6^ and cancer cell lines^7^. C2C12 myoblasts, the cell line most widely used to study myogenesis and myopathies, provide another relevant example: they rapidly lose their capacity to differentiate into myotubes upon passaging^8^. Third, overexpression of a ligase-fused bait can produce false positives^9–14^. For proteins that turn over slowly, the exogenous bait may fail to replace the long-lived, endogenous pool and localize elsewhere. Finally, even with TurboID – an engineered enzyme with faster kinetics – *in vivo* applications often require repeated or prolonged biotin exposure to achieve sufficient labeling^15–18^. Such incubation windows preclude capturing dynamic biological events and increase background. Extended exposure to high concentrations of biotin can also perturb cellular physiology^19^.

Here, we present interactome mapping methods for cultured cells and zebrafish that (1) are generalizable and require minimal optimization, (2) are applicable without genetic manipulation, and (3) achieve sub-hour temporal resolution. Central to SureCLIP (subcellular-resolution reversible crosslinker-mediated co-immunoprecipitation) and DrCLIP (*Danio rerio* reversible crosslinker-mediated co-immunoprecipitation) are cleavable chemical crosslinkers, dithiobis(succinimidyl propionate) (DSP) and dimethyl 3,3′-dithiobispropionimidate (DTBP), which covalently but reversibly stabilize PPIs within their native contexts. We first demonstrate the utility of SureCLIP by mapping the interactomes of a cytoplasmic and a nuclear protein, Hic-5 and lamin B1. We then apply DrCLIP, a protocol adapted for zebrafish, to map the binding partners of kindlin-2 at 1, 3, and 5 days post-fertilization (dpf). DrCLIP-guided analysis revealed that this focal adhesion protein is essential for somite boundary formation. Together, SureCLIP and DrCLIP enable genetically unperturbed interactome mapping with sub-hour temporal resolution in cell culture and a vertebrate model, overcoming limitations associated with proximity labeling.

## RESULTS

### Establishing SureCLIP protocol

Although co-IP/MS is challenging to optimize for robust and specific interactor recovery, it retains one key advantage over proximity labeling: it is a purely biochemical approach that permits the use of cells and organisms “as-is.” We surmised that DSP, an amine-reactive crosslinker with a reducing-agent-cleavable spacer (**Fig. 1a**), could address its shortcomings and establish a next-generation PPI mapping method. By covalently tethering bait and prey proteins before lysis, DSP should enable stringent extraction and washes. Separately, if DSP could be cleaved under conditions that preserve the immunocomplex, it could enable antibody- and bait-free elution of interactors (**Fig. 1b**). IgG chains and bait are dominant contaminants that hinder mass spectrometry-based prey identification^20^. We chose Hic-5 (TGFB1I1)^21,22^, a well-established component of the focal adhesion complex, as a model bait (**Fig. S1a**). Its interactome has been characterized by both co-IP/MS and proximity labeling using tagged Hic-5^23,24^, providing reference datasets against which to evaluate SureCLIP.

**Figure 1.**
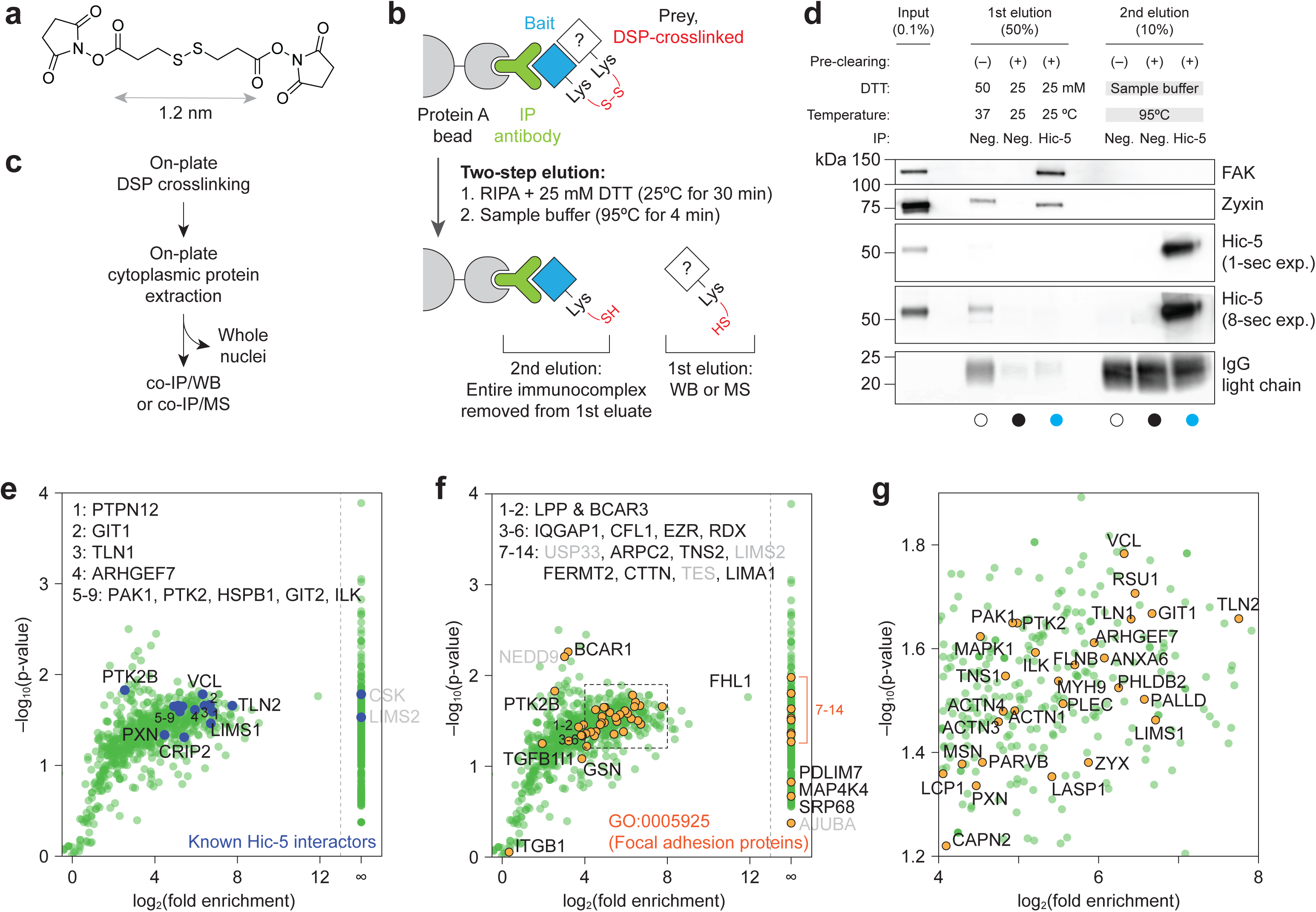
Proteome-wide identification of Hic-5 interactors by SureCLIP. **(a)** Chemical structure of DSP. **(b)** Schematic representation of the two-step SureCLIP elution protocol. **(c)** SureCLIP workflow for cytoplasmic proteins. WB: western blotting. **(d)** Immunoblots comparing the levels of the bait (Hic-5), IgG light chain, and preys (FAK and zyxin) in the input, first eluate, and second eluate from Hic-5 SureCLIP. Exp.: exposure; Neg.: negative (IgG isotype control). Hollow, blue, or black circles denote matching first- and second-elution pairs. (**e, f, and g**) Mass spectrometry analysis of the first elution fraction from Hic-5 SureCLIP. Green, all detected proteins; gray labels, proteins identified by a single tryptic peptide. **(e)** Navy, known Hic-5 interactors annotated in UniProt (Q62219). (**f** and **g**) Orange, focal adhesion proteins (GO:0005925). (**g**) Zoomed-in view of the dotted region in panel f.

As the first step, we crosslinked cells directly on the plate using DSP, covalently stabilizing PPIs as closely as possible to their native state (**Fig. 1c**). We then established an extraction protocol that enriches for cytoplasmic proteins from DSP-treated cells (**Fig. S1b**). Next, we asked whether this lysate could serve as an input for Hic-5 co-IP. Our goal was to selectively release crosslinked prey into the “1st eluate” while retaining the bait and IgG on the matrix for the “2nd eluate” (**Fig. 1b**). We sought to achieve this without covalent antibody immobilization, which prevents IgG contamination but can compromise antibody specificity/affinity and promote bead aggregation. By incubating in 25 mM dithiothreitol (DTT) at 25°C, we successfully isolated the known Hic-5 interactors FAK and zyxin with minimal leaching of bait and IgG heavy and light chains (**Figs. 1d** and **S2**).

### Proteome-wide identification of Hic-5 interactors by SureCLIP

We subsequently analyzed the DTT eluates by mass spectrometry. We reasoned that three features of SureCLIP could yield datasets of comparable or potentially greater quality than those obtained by proximity labeling: (1) covalent DSP linkage allows lysis and wash conditions similar to those used in streptavidin/biotin pulldown; (2) our samples contain minimal bait, whereas self-biotinylation leads to bait co-purification in enzymatic methods; and (3) our elution method does not involve boiling, potentially limiting the recovery of non-specifically bead-associated proteins that survive washes (see below). SureCLIP identified 1,021 proteins by ≥2 peptides (**Supplementary Table 1**), comparable to typical BioID or TurboID coverage. More importantly, all but one of the previously reported Hic-5 interactors that we detected showed >20-fold enrichment, highlighting the specificity of our approach (**Fig. 1e**; blue data points). Known interactors that were not detected generally exhibited low RNA abundance, suggesting they may be absent at the protein level (**Fig. S3a**), or are nuclear proteins that are minimally present in the cytoplasmic input.

Our dataset included 52 Gene Ontology (GO)-annotated focal adhesion proteins, 40 of which exhibited >16-fold enrichment (**Figs. 1f-g**). Three interactors from this group, talin-1, vinculin, and kindlin-2, were individually validated via immunoblotting (**Fig. S3b**). To distinguish strong from weak interactors, data point size was scaled by abundance measured by mass spectrometry (**Fig. 2a**). This representation identified vinculin (VCL), a core component of the force transduction layer in focal adhesions^25^, as the most prominent Hic-5 interactor. Strong interactors were heavily modified with DSP, with lysines bearing half-DSP scars (**Fig. S4a**). For example, 23 of 75 lysines in vinculin reacted with DSP, and in talin-1, 21 of 163 (**Fig. S4b**). In summary, we determined the Hic-5 interactome in cultured cells without genetic manipulation or specialized capture matrices (e.g., GFP-Trap or V5-Trap), and under conditions that minimize biochemical and physical perturbation. Notably, when standard co-IP was performed using the same antibodies and a widely used commercial kit, even the top interactor, vinculin, was not recovered although Hic-5, the bait, was still isolated (**Fig. S4c**).

**Figure 2.**
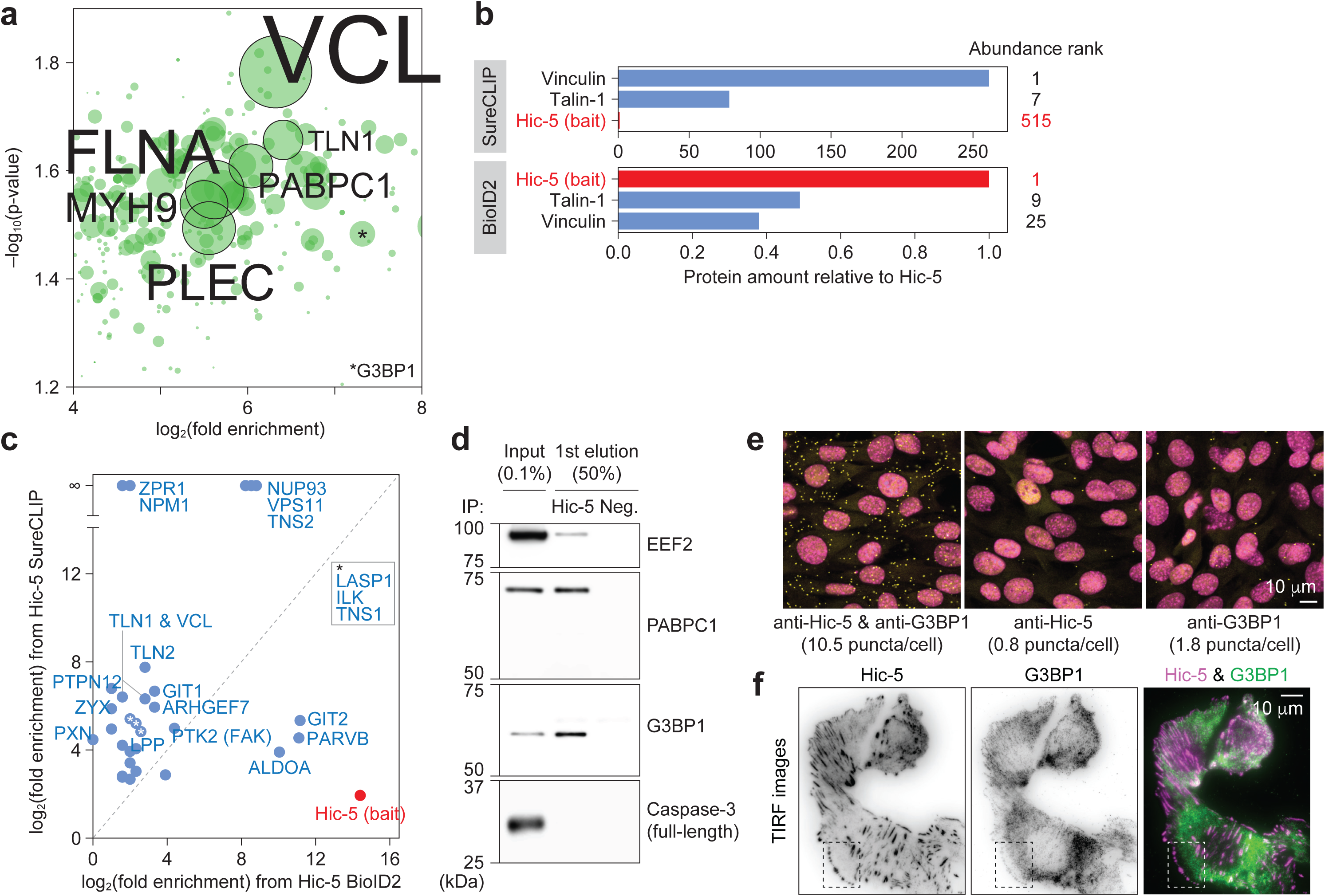
Prey-dominant elution in SureCLIP relative to BioID2 and validation of new Hic-5 interactors. **(a)** Mass spectrometry analysis of the first elution fraction from Hic-5 SureCLIP. Zoomed-in view of Fig. 1f. Data point and label sizes are scaled to protein abundance measured by mass spectrometry. The six most abundant proteins within this region are labeled. G3BP1, examined in panels **d-f**, is marked with an asterisk. **(b)** Relative abundance of proteins in the Hic-5 SureCLIP and BioID2 eluates, determined by mass spectrometry. **(c)** Fold enrichment comparison of proteins identified by Hic-5 SureCLIP and BioID2. Red, bait (Hic-5); blue, preys. **(d)** Immunoblot analysis of the input and first eluate from Hic-5 SureCLIP, probed for EEF2, PABPC1, G3BP1, and caspase-3. Caspase-3 was included as a non-interacting control. **(e)** PLA assessing the interaction between Hic-5 and G3BP1. Yellow puncta indicate PLA signals; nuclei in magenta. **(f)** TIRF images of C2C12 cells immunostained for Hic-5 and G3BP1.

### Comparison of Hic-5 interactome mapped by SureCLIP, BioID2, and conventional co-IP/MS

We compared our dataset with that of a recent study where Hic-5 interactors were identified using BioID2^24^. While differences in cell type (C2C12 versus U2OS), input material (cytoplasmic versus whole-cell lysates), methodology, and mass spectrometry instrumentation necessitate cautious interpretation, we identified three major differences.

First, the top hit from BioID2 was Hic-5 itself – an expected outcome of bait self-biotinylation (**Fig. 2b**). It showed 2.0- and 2.6-fold greater abundance than strong interactors such as talin-1 and vinculin. Hic-5 was likewise the top hit in conventional co-IP/MS^23^. In contrast, our eluate contained negligible amounts of Hic-5. It ranked as the ∼500^th^ most abundant protein and exhibited 78-fold and 260-fold lower abundance than talin-1 and vinculin, consistent with the immunoblotting results (**Fig. 1d**). The discrepancy between the two techniques was also reflected in the fold enrichment over control. BioID2 yielded a 22,000-fold enrichment of Hic-5, whereas SureCLIP yielded a 3.8-fold enrichment (**Fig. 2c**). Second, among 30 high-confidence hits in BioID2, 26 showed greater enrichment in our SureCLIP data (**Fig. 2c**). SureCLIP also identified more Hic-5 candidate interactors than BioID2, even under more stringent cutoffs (**Fig. S4d**). Lastly, we identified several interactors that support recent observations that focal adhesions harbor RNA-binding and translation-related proteins^26,27^. In addition to 20 focal adhesion proteins (GO:0005925, –log_10_(*p*) = 16.6), among our top 100 hits were 18 ribonucleoprotein granule proteins, 5 eukaryotic translation initiation factor 3 complex subunits, and 5 ribosomal proteins (GO:0035770, 0005852, and 0005840; –log_10_(*p*) = 15.3, 8.2, and 2.2) (**Fig. S4e** and **Supplementary Table 2**). None of these were meaningfully enriched by BioID2^24^ or conventional co-IP/MS^23^. Together, these findings demonstrate that SureCLIP matches or exceeds the capabilities of existing PPI mapping approaches while enabling the detection of new interaction partners.

### DTT-mediated elution improves dataset quality in SureCLIP

Our two-step elution protocol separates prey from bait and IgG. To better evaluate the extent to which this contributes to SureCLIP performance, we compared proteins recovered by DTT (“1st eluate”) with those released by directly boiling in sample buffer (“bulk eluate”) (**Supplementary Table 3**). All preceding steps were identical. Seven of the top 10 interactors were preserved across the two conditions, indicating that the elution strategy does not substantially affect the major hits (**Figs. S5a-b**), apart from the extreme enrichment of Hic-5 itself in bulk-eluted samples. However, a global comparison of fold enrichment across the two datasets revealed that most prey – including previously reported interactors and focal adhesion proteins in general – were more selectively recovered with DTT (**Fig. S5c**). Notably, protein abundance in the eluate obtained by boiling DTT-treated beads in sample buffer (“2^nd^ eluate”) strongly correlated with that in the bulk eluate (R^2^ = 0.997; **Fig. S6a**). The correlation between the DTT and bulk eluates was much lower (R^2^ = 0.499). This difference suggests that bulk elution recovers substantial amounts of non-specific material largely excluded by the stepwise approach. Consistent with this, the top 30 proteins more abundant in the bulk than in the DTT eluate generally showed poor enrichment (**Fig. S6b**).

### Hic-5-interacting RNA-binding proteins G3BP1 and PABPC1 show distinct colocalization patterns

We then used immunoblotting to individually validate three new Hic-5 interactors – G3BP1, PABPC1, and EEF2 – identified by SureCLIP (**Fig. 2d**). The interactions of Hic-5 with G3BP1 and PABPC1 were further supported by an orthogonal method, proximity ligation assay (PLA) (**Figs. 2e** and **S7a**). After confirming that the RNA-binding proteins G3BP1 and PABPC1 interact with Hic-5, we employed total internal reflection fluorescence (TIRF) microscopy to assess their spatial organization at or near the plasma membrane, where Hic-5 is primarily localized. Although G3BP1 is known to be diffusely distributed in the cytosol, TIRF imaging revealed its enrichment at focal adhesions in C2C12 myoblasts (**Fig. S7b**). We also observed colocalization of these proteins along the leading edge of lamellipodia (**Figs. 2f** and **S7c**). This aligns with the recent finding that G3BP1 knockdown decreases cell migration speed^27^. In contrast, PABPC1 showed no discernible localization at the lamellipodial rim (**Fig. S7d**) and did not appear to colocalize with Hic-5 near the cell surface. However, confocal imaging, which captures thicker z-sections than TIRF, revealed colocalization of Hic-5 and PABPC1 in bleb-like protrusions and filopodia-like structures (**Figs. S7e-f**). These observations suggest that, unlike G3BP1, PABPC1 is less directly involved in adhesion and instead may play a more prominent role in the nuclear export of Hic-5, similar to its reported function with the Hic-5 paralog paxillin^28^.

### Proteome-wide identification of lamin B1 interactors by SureCLIP

Having established SureCLIP for a cytoplasmic bait, we turned to the nucleus. Co-IP/MS is particularly challenging for nuclear envelope proteins because their extraction requires harsh detergents that disrupt PPIs. BioID was first developed in part for this reason, with early applications targeting components of the nuclear lamina, nuclear pore complex, and LINC (linker of nucleoskeleton and cytoskeleton) complex^3,29–31^. However, overexpression of these proteins can cause mislocalization and perturb chromatin organization, gene expression, nuclear morphology, and nucleocytoplasmic transport^10,32–35,13,14^. We thus asked whether SureCLIP could map the lamin B1 interactome without genetic manipulation.

We adapted the cytoplasmic extraction protocol to obtain a DSP-crosslinked nuclear lysate (**Figs. 1c**, **3a**, and **S8a**). Using an antibody against lamin B1, we co-purified lamin B2 without detectable contamination by fibrillarin or histone H2B (**Fig. 3b**). Of note, lamin B1, the bait, was present in both the first and second eluates, reflecting its ability to self-oligomerize^36^ (**Figs. 3b-c**). The amount of lamin B1 recovered, however, was comparable to that in 0.1% of the input, indicating relatively low pulldown efficiency. We found that sonication during nuclear extraction was essential for efficient capture (**Fig. 3b**), and analyzed the resulting DTT eluates by mass spectrometry.

**Figure 3.**
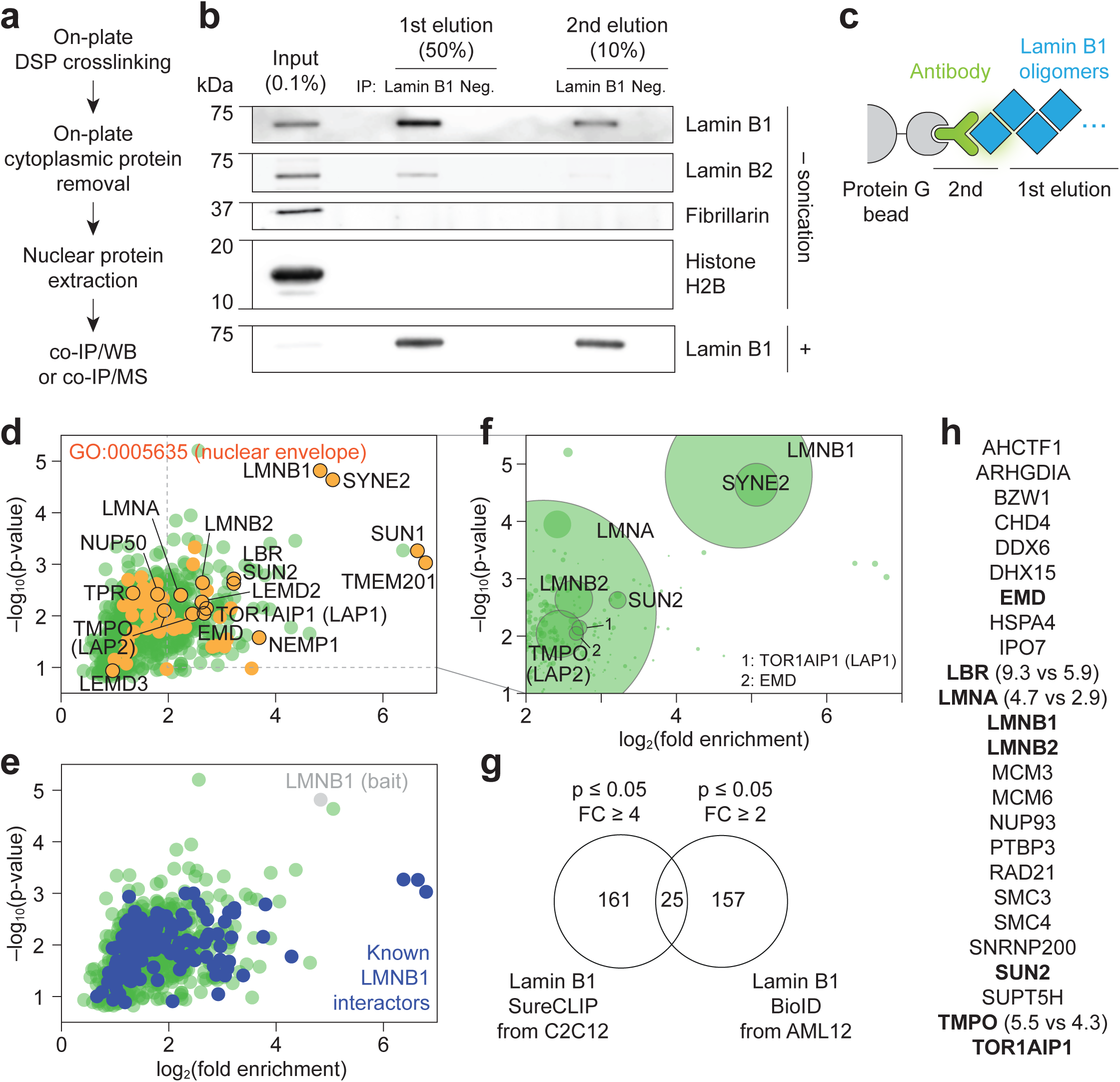
Proteome-wide identification of lamin B1 interactors by SureCLIP. **(a)** SureCLIP workflow for nuclear proteins. **(b)** Immunoblot analysis of the input and eluates from lamin B1 SureCLIP, probed for lamin B1, lamin B2, fibrillarin, and histone H2B, with or without sonication. **(c)** Schematic of lamin B1 oligomers distributed across the first and second eluates. (**d**, **e**, and **f**) Mass spectrometry analysis of the first elution fractions from lamin B1 SureCLIP. Green, all detected proteins. (**d**) Orange, nuclear envelope proteins (GO:0005635). (**e**) Navy, known lamin B1 interactors annotated in BioGRID. (**f**) Zoomed-in view of the dashed region in panel **d**, with data point size scaled to protein abundance measured by mass spectrometry. **(g)** Overlap between lamin B1 interactors identified by SureCLIP and by BioID.**(h)** The 25 proteins shared between the two datasets in panel **g**. Bold, nuclear lamina and associated proteins. Parentheses, fold enrichment by SureCLIP versus BioID, for proteins with values in both datasets.

Using SureCLIP, we detected 145 previously reported lamin B1 interactors and 66 nuclear envelope proteins (**Figs. 3d-e** and **Supplementary Table 4**). Our dataset included nearly all key proteins known to span the nuclear lamina and inner nuclear envelope, such as SUN1/2, LAP1/2, lamin B receptor (LBR), and emerin (EMD). TMEM201 (also known as SAMP1), a protein essential for myogenesis^37^, was the most highly enriched prey in C2C12 myoblasts. By abundance, the top nuclear interactors of lamin B1 were lamin A/C and lamin B1 itself, consistent with the mesh-like architecture of the nuclear lamina, followed by LAP2, nesprin-2 (SYNE2), lamin B2, and SUN2 (**Fig. 3f**). The presence of LAP2 in the DTT eluate was validated by immunoblotting (**Fig. S8b**). Finally, we compared the lamin B1 SureCLIP dataset with published BioID results^38^. As with the Hic-5 comparison, differences in cell type, methodology, and instrumentation warrant cautious interpretation. Nevertheless, SureCLIP identified a similar number of prey proteins despite using a more stringent cutoff (4-fold versus 2-fold; **Fig. 3g**), and recovered core nuclear envelope proteins as did BioID (**Fig. 3h**, bold). Among proteins for which fold enrichment could be compared, SureCLIP consistently yielded higher enrichment. In sum, SureCLIP profiles interactomes of cytoplasmic and nuclear baits alike, without genetic manipulation and with coverage and enrichment comparable to or greater than proximity labeling.

### DSP enables interactome mapping in zebrafish embryos

Having validated SureCLIP across subcellular fractions, we sought to establish the zebrafish counterpart, DrCLIP. PPI analysis remains challenging in this vertebrate model. Both co-IP/MS^39–41^ and proximity labeling^15–18^ have largely relied on microinjecting hundreds to thousands of embryos with mRNA encoding a tagged bait or generating transgenic lines, which takes at least 6 months. Proximity labeling additionally requires 12-18 hours of reaction time, even with TurboID and near-saturating biotin (∼0.5 mM)^15–18^. A window this long cannot resolve embryogenesis or other processes that unfold within hours. Interactome mapping in zebrafish is thus constrained by labor, lead time, and temporal resolution.

We asked whether DSP could be leveraged to overcome these barriers. We first adapted a standard deyolking protocol for DSP-crosslinked embryos to increase relative bait abundance and reduce background (**Fig. S9a**)^42,43^. Non-reducing SDS-PAGE revealed that the lysate from overnight DSP-treated, deyolked embryos contained crosslinked proteins that failed to migrate out of the loading well, similar to those observed in DSP-treated C2C12 cells (**Fig. 4a**). To assess crosslinking kinetics, we incubated embryos in 1 mM DSP for 0, 1, 4, 8, or 20 hours, then resolved and Coomassie-stained the lysates (**Fig. S9b**). Signal below 50 kDa declined from 1 hour, signal above 250 kDa rose from 4 hours, and a band appeared at the bottom of the well by 8 hours. Macroscopically, embryo coiling and cardiac contractions ceased within 2 hours. These observations indicate that DSP captures PPIs with a temporal resolution on the order of hours.

**Figure 4.**
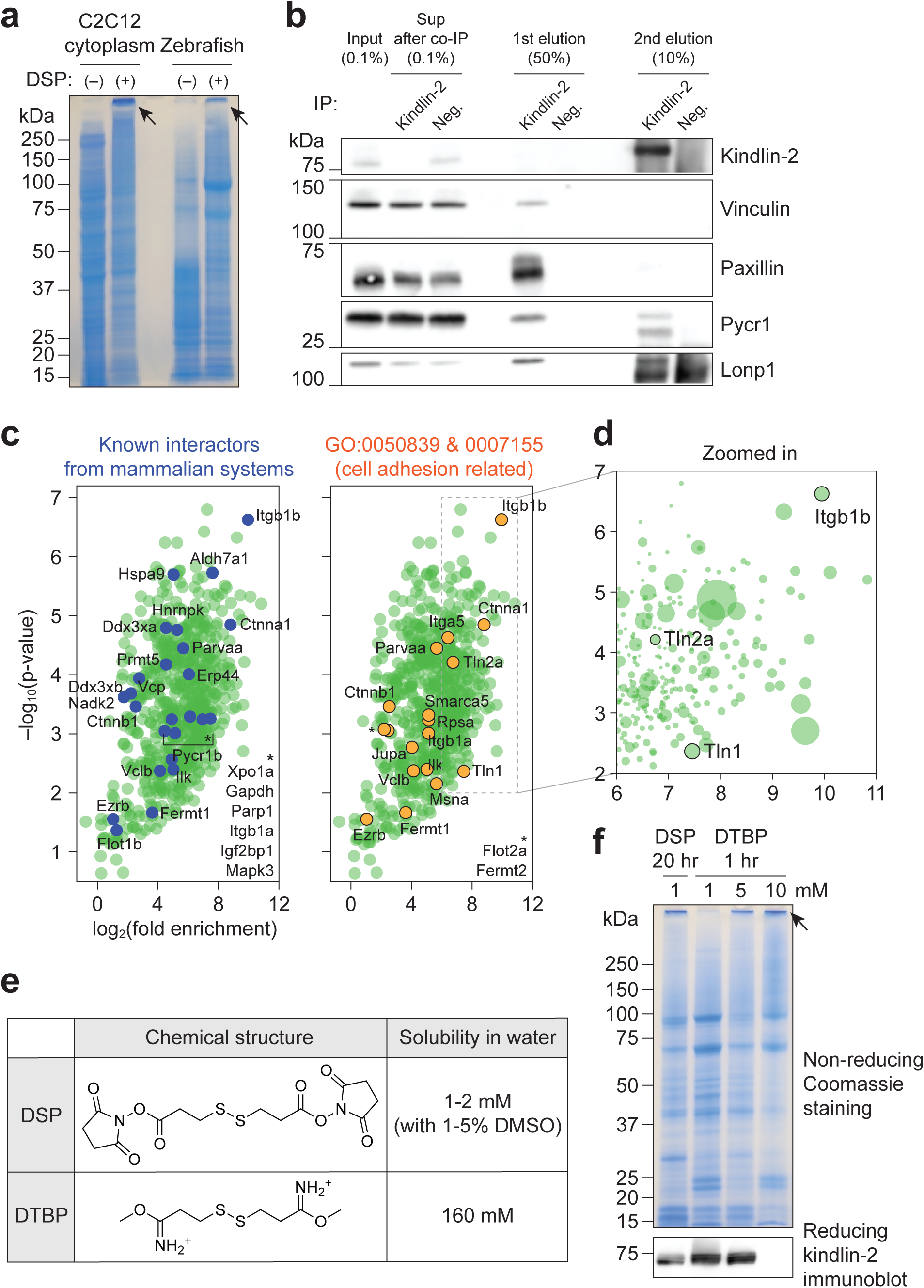
DSP enables interactome mapping in zebrafish embryos. **(a)** Coomassie staining of DSP-crosslinked and non-crosslinked lysates from C2C12 cells and deyolked 24-hpf zebrafish embryos. Arrows indicate proteins that failed to migrate out of the loading well. **(b)** Immunoblots comparing the levels of the bait (kindlin-2) and preys (vinculin, paxillin, Pycr1, and Lonp1) in the input, supernatant after co-IP, and first and second eluates from kindlin-2 DrCLIP. (**c** and **d**) Mass spectrometry analysis of the first elution fraction from kindlin-2 DrCLIP. Green, all detected proteins. (**c**) Left, navy, zebrafish orthologs of kindlin-2 interactors annotated in BioGRID from mammalian systems, plus Ddx3xa, Ddx3xb, Pycr1, and Prmt5. Right, orange, cell adhesion-related proteins (GO:0050839 and GO:0007155). (**d**) Zoomed-in view of the dashed region in the right panel of **c**, with data point size scaled to protein abundance measured by mass spectrometry. **(e)** Chemical structures and water solubility of DSP and DTBP. **(f)** Crosslinking of 3-dpf larvae with either DSP or DTBP. Lysates were analyzed by non-reducing SDS-PAGE and Coomassie staining (top), and by reducing immunoblotting for kindlin-2 (bottom). Arrow, crosslinked proteins retained in the loading well.

We then sought to isolate kindlin-2 (Fermt2) and its interactors from overnight-crosslinked embryos (**Fig. S9c**). Kindlin-2 is a focal adhesion protein that ranks among the most enriched at human myotendinous junctions (MTJs)^44^. Consistent with this, we found that zebrafish kindlin-2 localizes to somite boundaries, the future MTJ sites, at 24 hours post-fertilization (hpf; **Fig. S9d**). Two practical considerations made kindlin-2 a suitable model bait: (1) several of its interaction partners have been characterized in mammalian cells^45^ and (2) a commercial antibody raised against the human protein can immunoprecipitate, immunoblot, and immunostain zebrafish kindlin-2 due to 91.3% sequence identity (**Fig. S9e**). Protein A beads loaded with this antibody depleted kindlin-2 from lysates of DSP-crosslinked embryos (**Fig. 4b**). We eluted preys with DTT and detected paxillin and vinculin, two known interactors, by immunoblotting. Bimolecular fluorescence complementation independently confirmed both interactions (**Fig. S9f**). The DTT eluate also contained pyrroline-5-carboxylate reductase 1 (Pycr1), a mitochondrial matrix protein reported to interact with kindlin-2^46^, showing that our method can identify less prominent PPIs. Thus, the conditions established for DSP-crosslinked cells – immunoprecipitation, wash, and elution – proved equally effective for zebrafish. From non-crosslinked embryos, kindlin-2 was still fully immunoprecipitated, but none of the interactors were recovered (**Fig. S10a**), indicating that co-isolation of preys is DSP-dependent.

To profile the kindlin-2 interactome in 24-hpf embryos, we subjected the DTT eluates to mass spectrometry (**Supplementary Table 5**). Zebrafish frequently carry two paralogs of a mammalian gene (e.g., vinculin a and b; *vcla* and *vclb*). To cross-reference our hits with known mammalian interactors, we searched the BioGRID database using each zebrafish gene name with and without the a/b suffix, and identified 21 proteins with direct or indirect evidence of kindlin-2 interaction (**Fig. 4c**). Furthermore, we found Ddx3x (a and b), Pycr1, and Prmt5, four recently reported interactors absent from BioGRID^46–48^ The top interactor by fold enrichment and significance was integrin β1 (Itgb1b), in line with the best-characterized role of kindlin-2 as an integrin activator^49–52^. Talins, which bind the intracellular domain of β integrins alongside kindlins, were also highly enriched (Tln1, 180-fold; Tln2a, 110-fold). The recovery of these and other adhesion proteins confirmed that the pulldown worked as intended. However, unlike the major interactors in Hic-5 and lamin B1 SureCLIP (**Figs. 2a** and **3f**), Itgb1b, Tln1, and Tln2a did not stand out when data points were scaled by abundance (**Fig. 4d**). We hypothesized that prolonged DSP treatment captured bystanders alongside true interactors. We therefore sought to shorten the crosslinking time, both to reduce this background and to improve temporal resolution.

### DTBP enables rapid, stage-resolved mapping of the kindlin-2 interactome

DTBP is a crosslinker functionally analogous to DSP – amine-reactive, with the same DTT-cleavable spacer – but ∼100-fold more water-soluble (160 versus 1-2 mM; **Fig. 4e**). We examined whether this greater solubility could be exploited for faster PPI capture. In 3-dpf larvae, a 1-hour treatment with 5 or 10 mM DTBP indeed crosslinked proteins to an extent comparable to a 20-hour treatment with 1 mM DSP (**Fig. 4f**). DTBP also acted faster physiologically. Twitching and heartbeat ceased within ∼15 minutes in 5 mM DTBP (**Supplementary Video 1**), versus ∼2 hours in 1 mM DSP. At 10 mM DTBP, however, kindlin-2 immunoreactivity was lost, likely because extensive lysine modification masked the epitope (**Fig. 4f**). At 5 mM, kindlin-2 was immunoprecipitated, and vinculin, paxillin, and Pycr1 co-purified as they had from DSP-crosslinked embryos (**Fig. S10b**). We therefore used 6 mM for 1 hour for subsequent experiments, close to the validated 5 mM condition and well below the concentration that abolished antibody binding.

We repeated kindlin-2 DrCLIP with DTBP at 1 dpf and compared the interactome with that obtained using DSP. DTBP yielded 258 proteins identified by two or more peptides, about one-third of the 768 obtained with DSP. Of the 258, 199 (77%) were shared with DSP, indicating that the DTBP interactome was largely a subset (**Figs. 5a-b**). Additionally, we performed DTBP-based DrCLIP at 3 and 5 dpf to test whether the method extends to later stages. At all time points, Itgb1b and Tln1 were among the most abundant high-confidence hits (>16-fold enrichment, p < 0.01; **Fig. 5c** and **Supplementary Table 6**). Other abundant hits included cohesin/condensin subunits (Smc1al and Smc2-4) and embryonic hemoglobins (Hbae and Hbbe families), which are highly expressed at these stages (**Fig. S11**) and likely represent background. In support of this, their recovery paralleled their stage-specific transcription: cohesin/condensin proteins were most prominent at 1 dpf and hemoglobins at 3 and 5 dpf (**Figs. 5c** and **S11**; *smc1al* and *hbbe1.3* shown as representatives). Furthermore, the former are predominantly nuclear and the latter predominantly erythroid, whereas kindlin-2 is cytoplasmic and enriched in muscle. Despite these proteins, integrin signaling (dre04518) was the most enriched Kyoto Encyclopedia of Genes and Genomes (KEGG) pathway among the high-confidence hits (**Figs. 5d-e**). This demonstrates that DrCLIP can capture functionally relevant interactors from embryonic and larval zebrafish.

**Figure 5.**
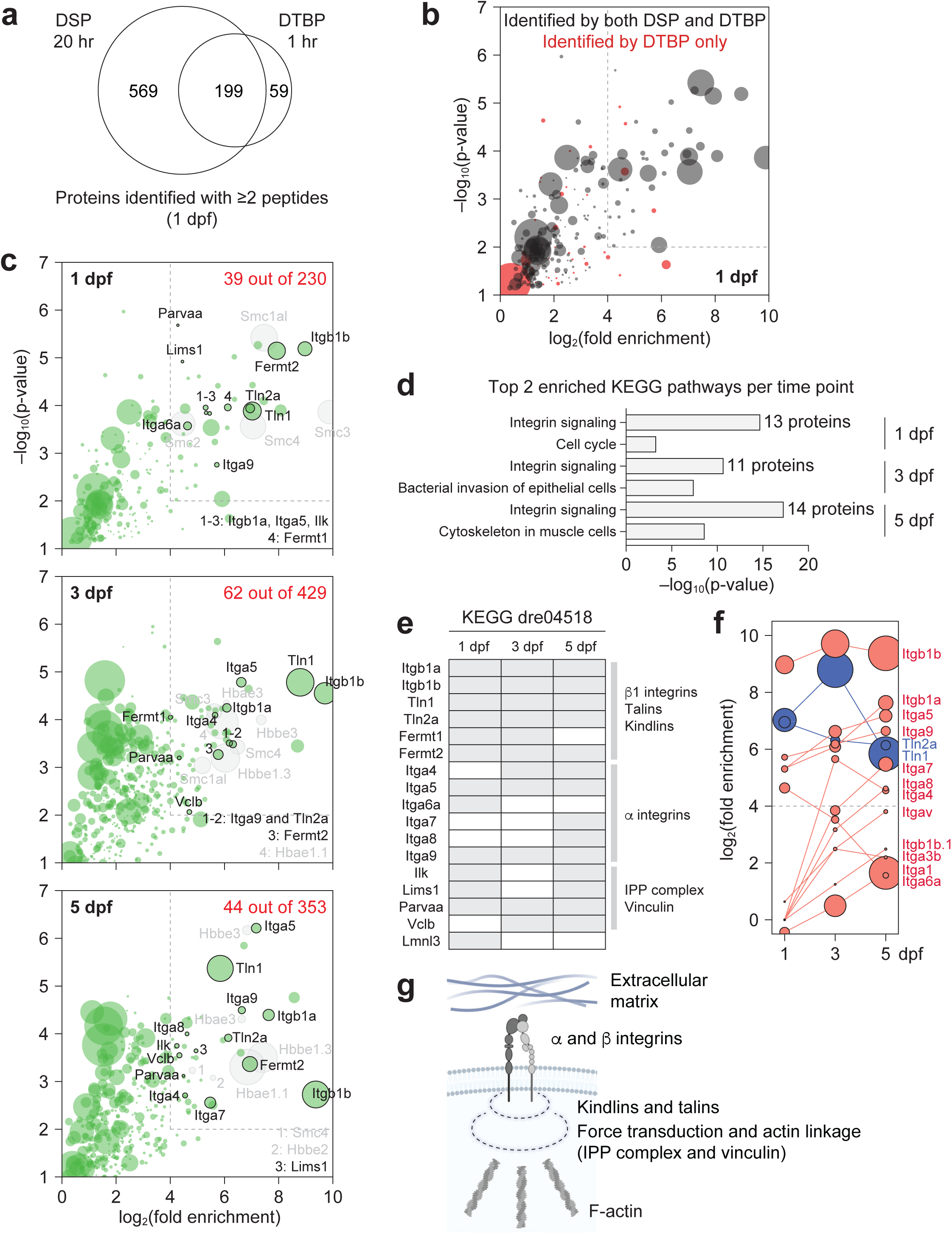
DTBP enables rapid, stage-resolved mapping of the kindlin-2 interactome. **(a)** Overlap between proteins identified by kindlin-2 DrCLIP using DSP (1 mM, 20 hours) or DTBP (6 mM, 1 hour) at 1 dpf. Contaminants (e.g., keratins) and vitellogenins (yolk proteins) were excluded. **(b)** Mass spectrometry analysis of the first elution fraction from DTBP-mediated kindlin-2 DrCLIP at 1 dpf. Gray, proteins also identified with DSP; red, proteins identified with DTBP only. Data point size is scaled to protein abundance. **(c)** Mass spectrometry analysis of the first elution fractions from DTBP-mediated kindlin-2 DrCLIP at 1, 3, and 5 dpf. Data point size is scaled to protein abundance. Dashed lines mark the high-confidence threshold (>16-fold enrichment, p < 0.01). Red numbers (“x out of y”) indicate high-confidence hits out of all proteins identified by ≥2 peptides with fold enrichment > 1. Gray labels, abundant proteins likely captured as background (cohesin/condensin subunits and embryonic hemoglobins). **(d)** Top two KEGG pathways enriched among high-confidence hits at each time point. **(e)** High-confidence hits annotated in KEGG integrin signaling (dre04518) at each time point. **(f)** Fold enrichment of integrins (red) and talins (blue) across time points. Data point size is scaled to protein abundance. **(g)** Schematic model of kindlin-2-associated proteins in developing zebrafish. Created in BioRender.

Beyond Itgb1b and Tln1, high-confidence hits at one or more time points included their paralogs Itgb1a and Tln2a, α integrins, the ILK-PINCH-parvin (IPP) complex components (Ilk, Lims1, and Parvaa), vinculin (Vclb), and kindlin-2 (Fermt2) (**Fig. 5e**). Although species, sample type, and method differed, orthologs of several of these hits (TLN2, ITGB1, ITGA4, ITGA5, ITGA6, ILK, and PARVA) were also identified by kindlin-2 BioID in HEK293 cells^53^. In zebrafish, β1 integrins and talins appear most proximal to kindlin-2 based on fold enrichment and abundance (**Fig. 5f**), consistent with the current model of mammalian focal adhesion architecture^49–52^. The α integrins were likely captured through their heterodimerization with β1 integrins and thus represent indirect interactors. Detection of the IPP subunits agrees with (1) the co-precipitation of UNC-112 (kindlin) with PAT-4 (ILK), UNC-97 (PINCH), and PAT-6 (parvin) in *C. elegans*^54^ and (2) the requirement for kindlin-2 in IPP binding to β integrin tails in cultured fibroblasts^55^. The presence of vinculin accords with its established role as a core focal adhesion protein^50^. As with the self-assembling lamin B1 (**Figs. 3c** and **3f**), recovery of kindlin-2 raises the possibility of self-association, which has been reported *in vitro* and in cell culture^56–58^. Further investigation will be needed to test this directly in live zebrafish. Taken together, our DrCLIP data suggest that, in developing zebrafish, kindlin-2 and talins engage multiple α/β1 integrin heterodimers and connect to downstream focal adhesion components, notably the IPP complex and vinculin (**Fig. 5g**).

### Kindlin-2 knockdown disrupts ECM assembly at somite boundaries and MTJ formation

α5β1 and α7β1 integrins mediate assembly of the extracellular matrix (ECM) at somite boundaries and anchor muscle cells to it^59–61^. Because kindlin-2 localized there (**Fig. S9d**) and associated with both integrins (**Fig. 5f**), we asked whether it is required for boundary formation. We first validated a previously published kindlin-2 morpholino^62^, confirming depletion at the protein level (**Fig. 6a**). At 2 dpf, morphants recapitulated the reported phenotypes: pericardial edema and bent, shortened trunks (**Fig. 6b**). At 3 dpf, the heartbeat persisted but morphants showed little or no spontaneous movement (**Supplementary Video 2**). We assessed myofiber organization and MTJ integrity at this stage by immunostaining for myosin heavy chain and vinculin, an established larval MTJ marker^61,63^. In morphants, MTJs were largely absent and myofibers disorganized (**Fig. 6c**). Earlier, at 28 hpf, somite boundaries were absent, with misaligned myofibers and gaps in the trunk musculature (**Figs. 6d** and **S12a**). More posteriorly, however, somites still exhibited their characteristic chevron shape (**Fig. S12b**), indicating that initial somite segmentation was unaffected. We therefore examined ECM assembly at boundaries, which underlies their maturation into MTJs, and found markedly reduced laminin deposition at 24 hpf (**Fig. S12c**). Kindlin-2 is thus dispensable for segmentation but required for boundary ECM assembly, and thereby MTJ formation and myofiber organization (**Fig. 6e**). Consistent with kindlin-2 directly activating β1-containing integrins, this phenotype resembles that of *itga5* mutants, in which initial somite segmentation is unaffected but anterior boundaries are not maintained^60^. It stands apart from the myotome defects reported for the downstream focal adhesion components paxillin, vinculin, Ilk, and β-parvin, in which segmental boundaries are retained^61,64,65^.

**Figure 6.**
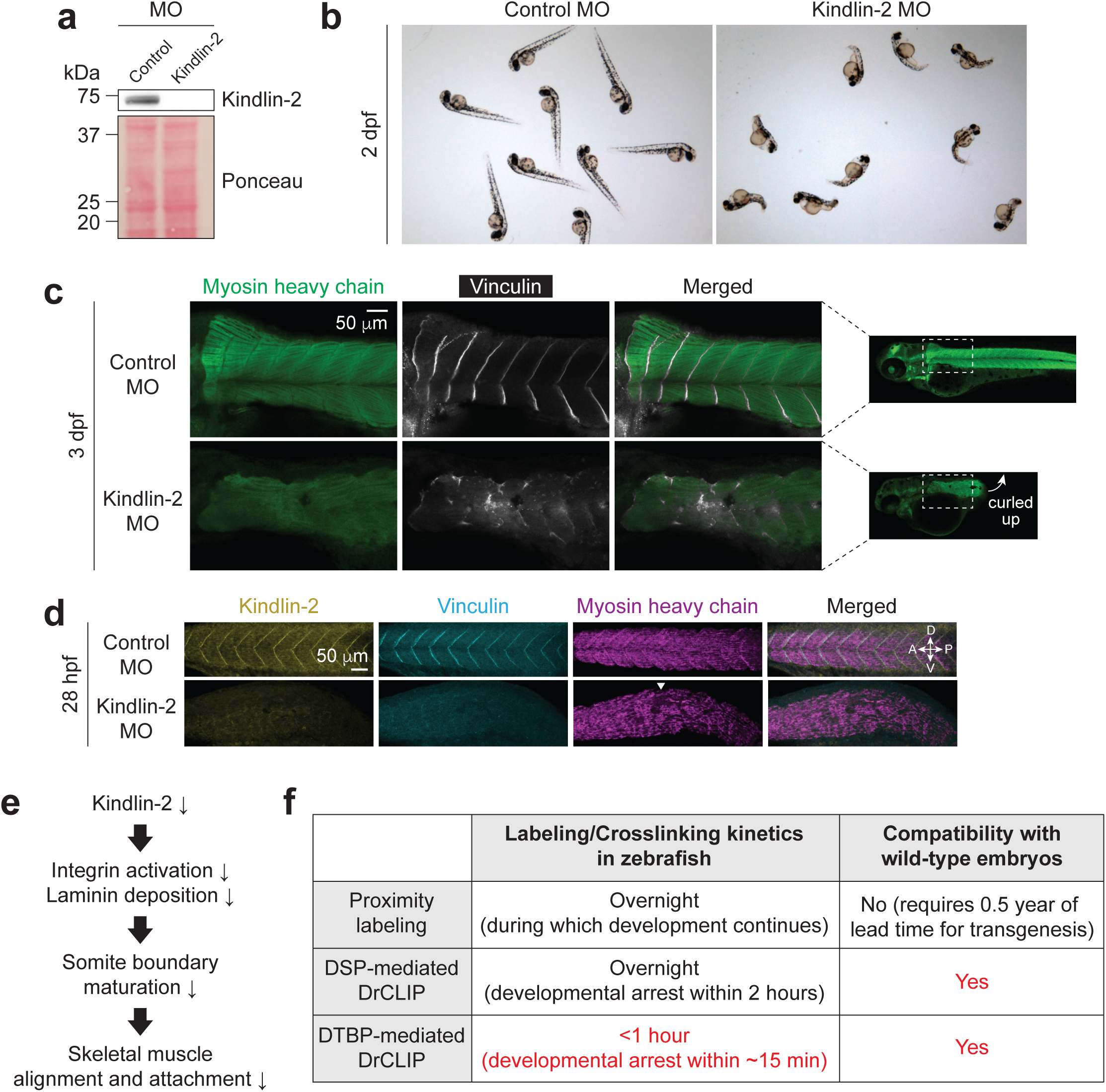
Kindlin-2 knockdown disrupts ECM assembly at somite boundaries and MTJ formation. **(a)** Immunoblot of kindlin-2 in control and kindlin-2 morpholino (MO)-injected embryos at 2 dpf. Ponceau S staining, loading control. **(b)** Live imaging of control and kindlin-2 morphants at 2 dpf. **(c)** Immunostaining of myosin heavy chain and vinculin in the trunk of control and kindlin-2 morphants at 3 dpf. Insets show the imaged region (dashed box). **(d)** Immunostaining of kindlin-2, vinculin, and myosin heavy chain in control and kindlin-2 morphants at 28 hpf. Arrowhead, gap in the trunk musculature. A, anterior; P, posterior; D, dorsal; V, ventral. **(e)** Model of kindlin-2 function at somite boundaries. **(f)** Comparison of proximity labeling and DrCLIP in zebrafish.

## DISCUSSION

DSP-mediated SureCLIP and DTBP-mediated DrCLIP enable protein interactome mapping in cells and zebrafish without genetic manipulation. By eliminating the need for transgenes, both methods avoid the associated lead time, labor, overexpression artifacts, and genetic drift. The capture window is comparable to that of proximity labeling in cultured cells (<30 minutes) and at least tenfold shorter in zebrafish (1 hour versus overnight; **Fig. 6f**). Overnight biotin labeling averages interactions across developmental stages and accumulates background throughout, whereas DTBP crosslinking arrests development within ∼15 minutes and progressively fixes PPIs. At 6 mM, the crosslinking profile showed little further change beyond the first 30 minutes (**Fig. S13a**), consistent with the ∼30-minute hydrolytic half-life of aliphatic imidoesters at pH 7-9^66^. The effective capture window for DrCLIP is thus shorter than the nominal 1-hour incubation.

Previous DSP-based co-IP/MS in cultured cells recovered <20 proteins, required covalent antibody-bead coupling that can compromise antibody affinity and specificity, and was incompatible with subcellular fractionation^67^. Combined with current-generation mass spectrometry, SureCLIP yielded several hundred protein identifications. Compared with published proximity labeling reference datasets, SureCLIP achieved greater fold enrichment. We discovered interactions between Hic-5 and the RNA-binding proteins G3BP1 and PABPC1, supported by PLA and colocalization imaging. For lamin B1, SureCLIP resolved the relative abundance of nuclear lamina proteins while avoiding the use of tagged lamins, which can mislocalize and alter nuclear morphology/mechanics^13,14^.

DrCLIP establishes a practical framework for endogenous interactome mapping in developing zebrafish. While DTBP has previously been used in this model organism^41^, prior work relied on a microinjected exogenous bait, a 5-hour incubation, and a buffer below the pH optimum for imidoester crosslinking. Addition of DTBP dihydrochloride, which we measured to have a pKa_1_ of 5.7 (see **Materials and Methods**; **Fig. S13b**), acidified this buffer further. To our knowledge, endogenous-bait PPI analysis in wild-type zebrafish has been reported in only two studies, both from the same group, using adult ovary and 1-cell stage embryos^68,69^. DrCLIP opens this capability to 1-5 dpf larvae, the stages most widely used in zebrafish research, with a 1-hour capture window. Applied to kindlin-2, DrCLIP recovered integrin complex components from 1 dpf onward. We therefore knocked down kindlin-2 and observed impaired somite boundary formation, followed by disruption of MTJ formation and myofiber alignment.

Several adaptations of conventional co-IP/MS have been introduced to improve robustness and specificity, each with limitations. For example, the two-step co-IP leverages biotinylated primary antibodies for sequential immunoprecipitations to reduce non-specific binders^70^. However, this method doubles the experimental time and does not resolve bait co-purification. Biotinylated antibodies are also not always commercially available. Quantitative multiplexed, rapid immunoprecipitation mass spectrometry of endogenous proteins (qPLEX-RIME) employs formaldehyde and disuccinimidyl glutarate, non-cleavable crosslinkers^71^. Like two-step co-IP, it does not eliminate bait contamination. Across four qPLEX-RIME baits, most high-confidence interactors showed a 2- to 16-fold enrichment, whereas two-step co-IP (two baits) and SureCLIP/DrCLIP (three baits) yielded enrichments ranging from 4- to 1,024-fold. We speculate that this reflects excessive crosslinking in qPLEX-RIME, which could increase background and contribute to a greater number of identifications (1,000-3,000 versus 250-1,000 for SureCLIP/DrCLIP).

Although our approach requires co-IP-compatible antibodies, this limitation is becoming less restrictive as endogenously tagged cell and zebrafish lines allow the use of anti-tag antibodies instead. Such lines are increasingly available from individual laboratories^72,73^ and through coordinated efforts (e.g., the Allen Institute’s collection of fluorescently tagged human iPSC lines and NIH-funded project R24OD040049). Lysine-free tags, such as HA or ALFA, are preferable; however, our analysis with kindlin-2 suggests that a suitable DSP/DTBP concentration range can preserve immunoreactivity while enabling sufficient crosslinking. Thus, lysine-containing tags, such as GFP or FLAG, may also be compatible with these methods.

The simplicity of SureCLIP and DrCLIP should facilitate broad adoption. The only non-standard reagents, DSP and DTBP, cost <$1 per replicate. DrCLIP may be adaptable to other model organisms, such as *C. elegans*, *D. melanogaster*, and *N. furzeri*. Because crosslinking occurs by immersion, the method may also apply to acute tissue slices and organoids, while perfusion could extend crosslinking to intact rodent tissues. Beyond live samples, extension to frozen murine and human tissues is an important next step. If successful, this would enable PPI analysis from biobank samples and support studies of disease-associated interactome changes.

## MATERIALS AND METHODS

Key reagents are listed in **Supplementary Table 7**. Mass spectrometry data have been deposited to ProteomeXchange via PRIDE (PXD071063).

### Cell culture

C2C12 cells, obtained from ATCC, were cultured in DMEM supplemented with 20% (v/v) FBS and 1% (v/v) penicillin-streptomycin, and were maintained at 37°C and 5% CO_2_. DMEM, penicillin-streptomycin, and FBS were purchased from Corning, Genesee Scientific, and Avantor, respectively (catalog #10-013-CV, 25-512, and 76419-584).

### Preparation of cytoplasmic lysate for SureCLIP

80-90% confluent 15-cm cell culture plates of C2C12 cells were washed three times with 5 mL of PBS. Following washes, 4 mL of crosslinking solution (0.5 mM DSP in PBS) was added to each dish and incubated with gentle orbital shaking for 25 min at room temperature. Additionally, dishes were manually tilted twice during this process to ensure the solution covered the entire plate. The DSP solution was then aspirated and the plate was washed with TBS. Cytoplasmic proteins were extracted with 1 mL of lysis buffer per plate (TBS supplemented with 1% NP40 (v/v) and 0.1% (v/v) SDS) for 25 min at room temperature on an orbital shaker. Successful extraction of cytoplasmic proteins can be monitored with a microscope; the plasma membrane should be removed while nuclei remain adherent and visible after extraction. Lysates were centrifuged at 600 x g for 5 min and the supernatant was transferred to a fresh tube and added protease inhibitor cocktail (Thermo Scientific #87785). Lysates were immediately used or stored at -80°C for long-term storage.

### Preparation of nuclear pellet for SureCLIP

DSP crosslinking and all steps through cytoplasmic extraction were performed as described above. Following collection of the cytoplasmic lysate, plates were washed with 5 mL TBS containing 0.01% NP-40. Nuclei were then harvested by adding 3 mL TBS + 0.01% NP-40 to each dish and gently scraping with a cell lifter (Corning #3008). The collected material was transferred to a 15- or 50-mL tube, and the plate was rinsed with an additional 10 mL of TBS + 0.01% NP-40; the rinse was combined with the initial suspension, yielding ∼13 mL per plate. Nuclei were centrifuged at 600 x g for 5 min, and the supernatant was carefully removed. The resulting DSP-crosslinked nuclear pellet was stored at -80°C until use. Typically, the pellet volume from a near-confluent 15-cm plate was ∼50 µL.

### Preparation of nuclear lysate for SureCLIP

For lysis, the frozen nuclear pellet was resuspended in four pellet volumes of RIPA buffer containing 0.7% SDS (higher than the standard 0.1%) and supplemented with protease inhibitor cocktail. After a 5-min incubation at room temperature, the gelatinous pellet and the liquid were transferred to a fresh 1.5 mL tube, rotated end-over-end for 10 min, gently pipetted up and down with a widened P1000 tip, rotated an additional 5 min, and passed ten times through an 18-gauge needle fitted to a 1 mL syringe. Following another 15 min of rotation, MgCl_2_ was added to 2 mM, mixed thoroughly, and Pierce Universal Nuclease was added to 500 U/mL. The mixture was rotated end-over-end for 20 min to reduce viscosity. Lysates were then sonicated in a Branson 1510 ultrasonic cleaner and clarified by centrifugation at 16,000 x g for 20 min at 4°C. The supernatant was transferred to a fresh tube, and protein concentration was determined by BCA assay (typically 1.5-2 mg/mL). Six volumes of TBS + 1% NP-40 were added to reduce the SDS concentration to 0.1% for antibody compatibility. Lysates were stored at -80°C until use.

### Zebrafish DSP crosslinking, deyolking, and protein extraction

*D. rerio* embryos (AB strain) were raised at 28.5°C in the UNC Zebrafish Aquaculture Core in compliance with protocol 25-005.0. Following manual dechorionation, groups of ∼100 embryos were transferred into 2 mL microcentrifuge tubes. After removing E2 medium as much as possible, 1 mL of freshly prepared 1 mM DSP solution in PBS, supplemented with protease inhibitors (10 µM pepstatin A, 1 mM phenylmethylsulfonyl fluoride, and 15 µM E-64) was added. Tubes were placed horizontally and incubated overnight on an orbital shaker at the lowest speed.

For deyolking, we modified a previously reported protocol^42^. After removing excess DSP solution, embryos were washed three times with 1.0 mL of deyolking wash buffer (111.0 mM NaCl, 3.5 mM KCl, 5.4 mM CaCl_2_, and 10.0 mM Tris; neutral pH). The final wash solution was replaced with 1 mL of deyolking buffer (55.0 mM NaCl, 1.8 mM KCl, 2.7 mM CaCl_2_, and 1.25 mM NaHCO_3_). Yolk sacs were then mechanically disrupted by pipetting up and down several times with a P20 micropipette and a 20 µL tip. Tubes containing embryos were then placed on an orbital shaker at 1100 RPM for 5 min, followed by centrifugation at 300 x g for 30 seconds. Supernatant was then discarded and replaced with 1.0 mL of deyolking wash buffer, followed by shaking at 1100 RPM for 2 min and another round of low-speed centrifugation (300 x g for 30 seconds). This process was repeated one more time. Finally, removal of yolk was verified under a dissecting scope.

Zebrafish proteins were then extracted utilizing RIPA buffer supplemented with protease inhibitor cocktail. 1.5 µL/embryo of protease inhibitor-supplemented RIPA buffer was added and the tube was rotated end-over-end at 20 RPM for 1 hour at room temperature. Lysate was then centrifuged at 13,000 x g for 20 min at 4°C. Supernatant was carefully transferred to a fresh tube and stored at -80°C until use. Protein extraction yields are shown in **Fig. S13c**.

### Zebrafish DTBP crosslinking, deyolking, and protein extraction

*D. rerio* embryos (AB strain) were raised at 28.5°C in the UNC Zebrafish Aquaculture Core in compliance with protocol 25-005.0. Following manual dechorionation, groups of 50-100 embryos were transferred into 1.7 mL microcentrifuge tubes. After removing E2 medium as much as possible, 1 mL of freshly prepared 6 mM DTBP (Thermo #20665) solution in 0.2 M bicine buffer (pH 8.5, Thermo #J61632.AE), supplemented with protease inhibitors (10 µM pepstatin A, 1 mM phenylmethylsulfonyl fluoride, and 15 µM E-64) was added. Tubes were placed horizontally and incubated for one hour at room temperature on an orbital shaker. Following incubation, the DTBP solution was replaced with wash buffer (containing 10 mM Tris), and incubated on ice for 20 minutes prior to proceeding with deyolking. For the DTBP kinetics experiment (**Fig. S13a**), embryos were incubated in wash buffer containing 50 mM Tris for at least 20 minutes to quench residual DTBP.

Embryos were then deyolked as mentioned above. Zebrafish proteins were then extracted utilizing RIPA buffer supplemented with protease inhibitor cocktail. 1.5 µL or 4 µL/embryo of protease inhibitor-supplemented RIPA buffer was added based on developmental stage (1 dpf or 3-5 dpf, respectively) and the tube was rotated end-over-end at 20 RPM for 1 hour at 4°C. After 30 minutes of RIPA incubation, 3 and 5 dpf embryos were manually disrupted briefly on ice using a tissue homogenizer followed by an additional 30 minutes of end over end rotation at 4°C. Lysate was then centrifuged at 13,000 x g for 20 min at 4°C. Supernatant was carefully transferred to a fresh tube and stored at -80°C until use. Protein extraction yields are shown in **Fig. S13c**.

### DTBP pKa measurement by spectrophotometric titration

The acid dissociation constants of DTBP were determined by spectrophotometric titration using a Pion SiriusT3 automated titrator equipped with a double-junction Ag/AgCl pH electrode and a UV dip-probe spectrophotometer. Measurements were performed at 25.0-25.1°C in ionic-strength-adjusted water (0.15 M KCl) under an argon atmosphere to exclude atmospheric CO_2_. The pH electrode was calibrated before each series of measurements by a four-parameter procedure (titration of 0.15 M KCl water with standardized 0.5 M HCl and 0.5 M KOH). To minimize hydrolysis, the sample was prepared immediately before the titration. A 10 mM stock solution of DTBP in water (5 µL) was added to the sample vial, and the instrument added 0.15 M KCl (aq) to a total volume of 1.5 mL, giving a final concentration of approximately 33 µM. Three sequential titrations were performed, each in the upward direction from pH 2 to pH 12 with 0.5 M KOH. Each titration comprised ∼50 spectra (210-450 nm) and was completed within 5 minutes. The pKa values were calculated by target factor analysis of the multiwavelength absorbance data using the Pion Refinement Pro software. The analysis gave two pKa values, corresponding to three species (AH ^2+^, AH^+^ and A): pKa = 5.7 and pKa_2_ = 10.6 (25.1°C, I = 0.183 M). The values reported are from the first titration, which was performed on the freshly prepared sample. The second ionization lies near the upper end of the titrated pH range (2 to 12), so pKa_2_ is less precisely determined than pKa_1_. Potentiometric titration gave comparable values, pKa_1_ = 6.0 and pKa_2_ = 10.1.

### Co-IP

#### Bead preparation

Protein A or G magnetic beads (Invitrogen #10001D or #10003D) were washed three times with PBS containing 2% (w/v) bovine serum albumin (BSA) and 0.01% (w/v) Tween-20 (“PBS-B/T”). For each co-IP experiment (a pair of experimental and control co-IPs), two tubes containing 20 µL of beads were rotated overnight in 500 µL PBS-B/T at 4°C. Three additional tubes containing 40 µL of beads were rotated overnight in 300 µL PBS-B/T supplemented with 6 µg of isotype control antibody (Proteintech #30000-0-AP (rabbit) or #B900620 (mouse)). Finally, one tube containing 40 µL of beads was rotated overnight in 300 µL PBS-B/T with 6 µg of the bait antibody: Proteintech #10565-1-AP (Hic-5), #11453-1-AP (kindlin-2), or #66095-1-Ig (lamin B1).

#### Pre-clearing and co-IP

For zebrafish co-IP experiments, ∼1.1 mg of crosslinked lysate was used per condition, whereas 2 mg of cytoplasmic or nuclear lysate was used for C2C12 cell co-IPs. Lysates from both cells and zebrafish underwent two pre-clearing steps: first with magnetic beads coupled to isotype control antibody, and then with empty beads. Both steps were performed on a rotator at 4°C for 45-60 min. Pre-cleared lysates were subsequently transferred to beads incubated with either control or bait antibody and rotated at 4°C for 3-4 hours.

#### Washes & elution

After co-IP, beads were washed six times with 1 mL of RIPA buffer prior to elution (3, 3, 5, 5, 10, and 10 min). Following the fourth wash, beads were transferred to a fresh tube using the fifth wash. Prey proteins were eluted with 50 µL of RIPA buffer containing 25 mM DTT. Beads were incubated in this solution for 30 min at 25°C with vigorous shaking, and the supernatant was carefully transferred to a Protein LoBind tube (Eppendorf #022431081) and stored at -80°C. (For “bulk” elution in **Figs. S5** and **S6**, this DTT elution step was omitted, and beads proceeded directly to the second elution.) A second elution was performed by adding 50 µL of 5x Laemmli sample buffer supplemented with 100 mM DTT directly to the beads, followed by boiling at 95°C for 4 min. The resulting supernatant was transferred to a fresh tube and stored at -80°C.

### C2C12 immunofluorescence

C2C12 cells were grown on a chambered coverglass (Ibidi #80826). Cells were fixed in PBS containing 2% (w/v) paraformaldehyde for 10 min at room temperature and washed with PBS three times. Fixed cells were permeabilized and blocked in immunofluorescence buffer (PBS containing 0.1% Triton X-100, 0.02% SDS, and 10 mg/mL BSA) for 30 min. The cells were then incubated with primary antibodies diluted in immunofluorescence buffer at room temperature for 2 hours, washed three times with PBS, and incubated with the appropriate secondary antibodies diluted in immunofluorescence buffer at room temperature for 45 min. Finally, the cells were washed with PBS three times, and added VECTASHIELD antifade mounting medium with DAPI (Vector Laboratories #H-1200-10). A Leica SP8 or Stellaris 8 FALCON STED confocal microscope equipped with a 63x oil-immersion objective was used to acquire micrographs. TIRF imaging was conducted using an Olympus IX81 or a Nikon N-STORM microscope with a 60x objective.

### C2C12 PLA

C2C12 cells were seeded on a chambered coverglass (Lab-Tek #155411). The PLA was performed using a DuoLink In Situ Red Starter Kit (Sigma #DUO92101) according to the manufacturer’s instructions. Images were acquired using a Leica Stellaris 8 FALCON STED confocal microscope equipped with a 63x oil-immersion objective.

### Zebrafish immunofluorescence

Zebrafish embryos were manually dechorionated and fixed in 4% (w/v) paraformaldehyde for 90 min. Following fixation, embryos were washed four times in “PBX” (PBS supplemented with 0.5% (v/v) Triton X-100) and then placed in blocking solution (10 mM Tris, 250 mM NaCl, 10% (v/v) horse serum, 0.5% (w/v) BSA in PBX) for 90 min. Primary antibodies – kindlin-2 (Proteintech #11453-1-AP), vinculin (Sigma #V9131), myosin-4 (Invitrogen #14-6503-82), and laminin (Sigma #L9393) – were diluted in blocking solution (1:200, 1:200, 1:400, and 1:100, respectively) and incubated with embryos overnight at 4°C with gentle agitation. After incubation, embryos were washed with PBX and subjected to a second 90-min blocking step at room temperature. Secondary antibodies (Invitrogen #A21121, #A21144, and #A31573) were diluted 1:200 in blocking solution, and embryos were incubated in this solution for 150 min at room temperature with gentle agitation. Following incubation, embryos were washed extensively with PBX and stored in PBS at 4°C until imaging. Optically sectioned images were acquired using a Leica Stellaris 8 FALCON STED confocal microscope equipped with a 5x, 10x, and 20x dry objectives.

### Zebrafish brightfield images and videos

Images in **Fig. 6b** were acquired with a Zeiss Axio Zoom V16 stereomicroscope equipped with an Axiocam 503 color camera. **Supplementary Videos 1** and **2** were acquired with an Accu-Scope EXI-410 microscope equipped with an Excelis HD camera (30 frame per second) and a 10x dry objective.

### Zebrafish morpholino microinjection

All morpholinos were purchased from Gene Tools and diluted in nuclease-free water. Embryos were microinjected with 1.0 nL of either 0.2 mM control (CCTCTTACCT CAGTTACAATTTATA) or kindlin-2 ATG (GCATCCGTATACCGTCCAGCGCCAT) morpholino at the 1-cell stage.

### Bimolecular fluorescence complementation

The *fermt2*, *pxna,* and *vclb* coding sequences (NCBI Gene ID: 553051, 399546, and 100334859) were inserted into either the pCS2+-mVenus^1-155^-GGGS plasmid (Addgene #162610) or mCerulean^156-239^-GGGS plasmid (Addgene #162616) via In-Fusion cloning (Takara Bio #638956). The mMESSAGE mMACHINE SP6 Transcription Kit (Invitrogen #AM1340) was used to generate mVenus^1-155^-kindlin-2, mCerulean^156-239^-paxillin, mCerulean^156-239^-vinculin, mVenus^1-155^, and mCerulean^156-239^ mRNAs. mRNA cocktails were prepared in matching 600 nM concentrations and mixed in a 1:1 volume ratio prior to injection as previously described^74^. 1 nL of combined mRNA was injected into each embryo at the 1-cell stage. 24-hpf embryos were dechorionated and embedded in low melting point agarose molds (1% (w/v) agarose in E2 medium containing 0.0168% (w/v) tricaine) prior to imaging. Optically sectioned images were acquired using a Leica Stellaris 8 FALCON STED confocal microscope equipped with a 10x dry objective.

### SDS-PAGE gel staining

#### Coomassie staining

A total of 20 µg of protein (crosslinked or non-crosslinked) was mixed with 4x non-reducing Laemmli SDS sample buffer (Thermo Scientific #J63615.AC) to a final concentration of 1x. These samples were not boiled. Lysates were loaded onto 4-12% pre-cast Bis-Tris gels (Invitrogen #NW04122BOX) for electrophoresis. Subsequently, gels were washed three times with ultrapure water, and all excess water was removed before adding Coomassie stain (Bio-Rad #1610786). Gels were shaken for 1 hour at room temperature, then destained and washed with ultrapure water overnight prior to imaging.

**Silver staining** was performed using the Pierce Silver Stain Kit (Thermo Scientific #24612), following manufacturer’s instructions.

### Immunoblotting

A total of 10-30 µg of protein per lane was mixed with Laemmli sample buffer containing 100 mM DTT to a final concentration of 1x. For co-IP/WB, total protein was not quantified at every stage of the experiment, instead the amount of protein loaded was represented as a fraction of the total sample or elution volume. All samples were boiled at 95°C for 4 min and loaded on Bis-Tris gels (Invitrogen) for electrophoresis. Proteins were transferred to a nitrocellulose membrane and stained with Ponceau S (Thermo Scientific #A40000279) to confirm equal protein loading and successful transfer. Membranes were then washed with TBS containing 0.05% (w/v) Tween-20 (“TBST”) several times. The membrane was blocked with 5% (w/v) non-fat milk powder in TBST for 15-30 min at room temperature and subsequently incubated with primary antibodies at 4°C overnight. Primary antibody was removed with washes of TBST. Chemiluminescent detection was conducted using either ProSignal Pico or Femto (Genesee Scientific #20-300 or #20-302) after 45-min incubation with horseradish peroxidase-conjugated secondary antibodies at room temperature. For fluorescence-based detection, Alexa Fluor Plus 680- or 800-conjugated secondary antibodies (Invitrogen #A32802 or #A32789) were used. Western blot images were obtained using a KwikQuant Imager (Kindle Biosciences) or Odyssey CLx (LI-COR).

### Hic-5 SureCLIP mass spectrometry

This experiment was performed at the Salk Mass Spectrometry Core. Samples were precipitated by methanol/chloroform and redissolved in 8 M urea/100 mM TEAB, pH 8.5. Proteins were reduced with 5 mM TCEP hydrochloride and alkylated with 10 mM chloroacetamide. Proteins were digested overnight at 37°C in 2 M urea/100 mM TEAB, pH 8.5, with trypsin (Promega). The digested peptides were labeled with TMT, and pooled samples were fractionated by basic reversed phase (Thermo Scientific #84868).

The TMT labeled samples were analyzed on a Orbitrap Eclipse Tribrid mass spectrometer. Samples were injected directly onto a 25-cm, 100-μm ID column packed with BEH 1.7-μm C18 resin. Samples were then separated at a flow rate of 300 nL/min on an EASY-nLC 1200. Buffer A and B were 0.1% formic acid in water and 90% acetonitrile, respectively. A gradient of 1-15% B over 30 min, an increase to 45% B over 120 min, an increase to 100% B over 20 min and held at 100% B for 10 min was used for a 180 min total run time.

Peptides were eluted directly from the tip of the column and nanosprayed directly into the mass spectrometer by application of 2.5 kV voltage at the back of the column. The Eclipse was operated in a data dependent mode. Full MS1 scans were collected in the Orbitrap at 120k resolution. The cycle time was set to 3 seconds, and within this 3 seconds the most abundant ions per scan were selected for CID MS/MS in the ion trap. The TMT samples were analyzed by MS3 analysis with multinotch isolation (SPS3) was utilized for detection of TMT reporter ions at 60k resolution. Monoisotopic precursor selection was enabled and dynamic exclusion was used with exclusion duration of 60 seconds.

Protein and peptide identification were done with Integrated Proteomics Pipeline – IP2 (Integrated Proteomics Applications). Tandem mass spectra were extracted from raw files using RawConverter and searched with ProLuCID against UniProt mouse database. The search space included all fully-tryptic and half-tryptic peptide candidates. Carbamidomethylation on cysteine and TMT on lysine and peptide N-term were considered as static modifications. DSP samples included a differential modification of +145.0197245 Da at lysine to account for the remaining tag of the crosslinker. Data was searched with 50 ppm precursor ion tolerance and 600 ppm fragment ion tolerance. Identified proteins were filtered to using DTASelect and utilizing a target-decoy database search strategy to control the false discovery rate to 1% at the protein level. Quantitative analysis of TMT was done with Census filtering reporter ions with 10 ppm mass tolerance and 0.6 isobaric purity filter.

### Lamin B1 SureCLIP and kindlin-2 DrCLIP mass spectrometry

#### Sample preparation

Immunoprecipitated samples (DTT elutions from lamin B1 SureCLIP and kindlin-2 DrCLIP) were subjected to SDS-PAGE and stained with Coomassie. Lanes (1 cm) for each sample were excised, and the proteins were reduced with 10 mM DTT for 30 min at 56°C, alkylated with 15 mM iodoacetamide for 45 min in the dark at room temperature, and in-gel digested with trypsin (3 μg) overnight at 37°C. Peptides were extracted, desalted with C18 spin columns (Pierce) and dried via vacuum centrifugation. Peptide samples were stored at -80°C until further analysis.

#### LC/MS/MS analysis

Samples were analyzed in a randomized order by LC-MS/MS using a Vanquish Neo coupled to an Orbitrap Astral mass spectrometer (Thermo Scientific). A pooled sample was analyzed at the beginning and end of the sequence. Samples were injected onto an IonOpticks Aurora series 3 C18 column (75 μm ID × 15 cm, 1.6 μm particle size; IonOpticks) and separated over a 30-min method. The gradient for separation consisted of 2-30% mobile phase B at a 300 nL/min flow rate, where mobile phase A was 0.1% formic acid in water and mobile phase B consisted of 0.1% formic acid in 80% ACN. The Orbitrap Astral was operated in Data Dependent Acquisition (DDA) mode in which the most intense precursors were selected within a 0.6-second cycle for subsequent HCD fragmentation and Astral MS/MS detection. A full MS scan (m/z 375-1500) was collected; resolution was set to 180,000 with a maximum injection time of 5 ms and AGC target of 300%. MS/MS scans (m/z 110-2000) were acquired in the Astral analyzer with a maximum injection time of 2.5 ms and AGC target of 100%. The normalized collision energy was set to 30% for HCD, with an isolation window of 2 m/z.

#### Data analysis

Raw data files were processed in Proteome Discoverer (version 3.1, Thermo) and searched against the UniProt mouse (containing ∼17,000 entries) or zebrafish database (containing ∼27,000 entries), appended with the MaxQuant common contaminants database (∼300 entries). The following settings were used: enzyme specificity set to trypsin, up to two missed cleavages allowed, methionine oxidation, and N-terminal acetylation set as variable modifications. For lamin B1 SureCLIP, half-DSP (+87.9983 Da) and half-DSP + acetamide (+145.0197 Da) on lysine residues were also set as variable modifications; for kindlin-2 DrCLIP, half-DTBP neutral (+87.0143 Da), half-DTBP protonated (+88.0221 Da), and half-DTBP + acetamide protonated (+145.0436 Da). A false discovery rate (FDR) of 5%/1% protein/peptide was used to filter all data. Imputation from a normal distribution, and statistical analysis were performed in Perseus software (version 1.6.14.0). Proteins with log_2_ fold change of at least 1 and a q-value of less than 0.05 (Student’s t-test, FDR-permutation) were considered significant.

## COMPETING FINANCIAL INTERESTS

The authors declare no competing financial interests.

## ACKNOWLEDGMENTS

U.H.C. is grateful to Martin Hetzer (Institute of Science and Technology Austria), Tony Hunter (Salk Institute for Biological Studies), and Lorenzo Puri (Sanford-Burnham-Prebys) for their mentorship during postdoctoral training. We thank Jongmin Kim (Cornell University), Whitney Edwards (UNC), and Jongmin Park (Yonsei University) for critical reading of the manuscript. We also thank the staff members of the UNC Metabolomics and Proteomics, Zebrafish Aquaculture, Hooker Imaging, and Biology Microscopy core facilities for their technical support, especially Laura Herring, Angie Mordant, Michelle Altemara, Wendy Salmon, and Nathanaël Prunet. This work was supported by the National Institutes of Health grants K01AR080828 (to U.H.C.), R35GM130312 (to J.E.B.), 1S10OD030300 (to the Hooker Imaging Core), the National Science Foundation Graduate Research Fellowship DGE-2439854 (to S.A.A.), and 2P30CA016086-45 (to the Lineberger Comprehensive Cancer Center and Metabolomics and Proteomics Core Facility). Additional support was provided by UNC Chapel Hill start-up funding from the Lineberger Comprehensive Cancer Center and the Department of Cell Biology and Physiology (to U.H.C.).

## AUTHOR CONTRIBUTIONS

⍰ Conceptualization: U.H.C.
⍰ Investigation: B.T.C., C.D.T., W.B.F., S.A.A., M.H., H.L., and U.H.C.
⍰ Formal analysis and manuscript writing: B.T.C. and U.H.C.
⍰ Manuscript review and editing: all authors
⍰ Supervision: J.E.B. and U.H.C.

## Notes

### Competing Interest Statement

The authors have declared no competing interest.

### Summary of Updates

Added DrCLIP data generated with the crosslinker DTBP (mainly Figures 5 and 6).

## REFERENCES

1. Gingras, A.-C., Gstaiger, M., Raught, B. & Aebersold, R. Analysis of protein complexes using mass spectrometry. Nat. Rev. Mol. Cell Biol. 8, 645–654 (2007).

2. Dunham, W. H., Mullin, M. & Gingras, A.-C. Affinity purification coupled to mass spectrometry: basic principles and strategies. Proteomics 12, 1576–1590 (2012).

3. Roux, K. J., Kim, D. I., Raida, M. & Burke, B. A promiscuous biotin ligase fusion protein identifies proximal and interacting proteins in mammalian cells. J. Cell Biol. 196, 801–810 (2012).

4. Rhee, H.-W. et al. Proteomic mapping of mitochondria in living cells via spatially restricted enzymatic tagging. Science 339, 1328–1331 (2013).

5. Branon, T. C. et al. Efficient proximity labeling in living cells and organisms with TurboID. Nat. Biotechnol. 36, 880–887 (2018).

6. Selich, A. et al. Massive clonal selection and transiently contributing clones during expansion of mesenchymal stem cell cultures revealed by lentiviral RGB-barcode technology. Stem Cells Transl. Med. 5, 591–601 (2016).

7. Ben-David, U. et al. Genetic and transcriptional evolution alters cancer cell line drug response. Nature 560, 325–330 (2018).

8. Shintani-Ishida, K., Tsurumi, R. & Ikegaya, H. Decrease in the expression of muscle-specific miRNAs, miR-133a and miR-1, in myoblasts with replicative senescence. PLOS ONE 18, e0280527 (2023).

9. Nix, D. A. et al. Targeting of zyxin to sites of actin membrane interaction and to the nucleus. J. Biol. Chem. 276, 34759–34767 (2001).

10. Griffis, E. R., Xu, S. & Powers, M. A. Nup98 localizes to both nuclear and cytoplasmic sides of the nuclear pore and binds to two distinct nucleoporin subcomplexes. Mol. Biol. Cell 14, 600–610 (2003).

11. Kedersha, N. et al. Stress granules and processing bodies are dynamically linked sites of mRNP remodeling. J. Cell Biol. 169, 871–884 (2005).

12. Crisp, M. et al. Coupling of the nucleus and cytoplasm: role of the LINC complex. J. Cell Biol. 172, 41–53 (2006).

13. Kaneshiro, J. M., Capitanio, J. S. & Hetzer, M. W. Lamin B1 overexpression alters chromatin organization and gene expression. Nucleus 14, 2202548 (2023).

14. Odell, J. & Lammerding, J. N-terminal tags impair the ability of lamin A to provide structural support to the nucleus. J. Cell Sci. 137, jcs262207 (2024).

15. Pronobis, M. I., Zheng, S., Singh, S. P., Goldman, J. A. & Poss, K. D. In vivo proximity labeling identifies cardiomyocyte protein networks during zebrafish heart regeneration. eLife 10, e66079 (2021).

16. Rosenthal, S. M. et al. A toolbox for efficient proximity-dependent biotinylation in zebrafish embryos. Mol. Cell. Proteomics 20, 100128 (2021).

17. Xiong, Z. et al. In vivo proteomic mapping through GFP-directed proximity-dependent biotin labelling in zebrafish. eLife 10, e64631 (2021).

18. Tetenborg, S. et al. Uncovering the electrical synapse proteome in retinal neurons via in vivo proximity labeling. eLife 14, RP105935 (2025).

19. Griffin, J. B., Rodriguez-Melendez, R., Dode, L., Wuytack, F. & Zempleni, J. Biotin supplementation decreases the expression of the SERCA3 gene (ATP2A3) in Jurkat cells, thus, triggering unfolded protein response. J. Nutr. Biochem. 17, 272–281 (2006).

20. Antrobus, R. & Borner, G. H. H. Improved elution conditions for native co-immunoprecipitation. PLOS ONE 6, e18218 (2011).

21. Thomas, S. M., Hagel, M. & Turner, C. E. Characterization of a focal adhesion protein, Hic-5, that shares extensive homology with paxillin. J. Cell Sci. 112, 181–190 (1999).

22. Alpha, K. M., Xu, W. & Turner, C. E. Paxillin family of focal adhesion adaptor proteins and regulation of cancer cell invasion. Int. Rev. Cell Mol. Biol. 355, 1–52 (2020).

23. Petropoulos, C. et al. Roles of paxillin family members in adhesion and ECM degradation coupling at invadosomes. J. Cell Biol. 213, 585–599 (2016).

24. Brock, K. et al. A comparative analysis of paxillin and Hic-5 proximity interactomes. Cytoskeleton 82, 12–31 (2025).

25. Carisey, A. & Ballestrem, C. Vinculin, an adapter protein in control of cell adhesion signalling. Eur. J. Cell Biol. 90, 157–163 (2011).

26. Katz, Z. B. et al. Mapping translation ‘hot-spots’ in live cells by tracking single molecules of mRNA and ribosomes. eLife 5, e10415 (2016).

27. Boraas, L. C. et al. G3BP1 ribonucleoprotein complexes regulate focal adhesion protein mobility and cell migration. Cell Rep. 44, 115237 (2025).

28. Woods, A. J., Kantidakis, T., Sabe, H., Critchley, D. R. & Norman, J. C. Interaction of paxillin with poly(A)-binding protein 1 and its role in focal adhesion turnover and cell migration. Mol. Cell. Biol. 25, 3763–3773 (2005).

29. Kim, D. I., et al. Probing nuclear pore complex architecture with proximity-dependent biotinylation. Proc. Natl. Acad. Sci. USA 111, E2453–E2461 (2014).

30. Kim, D. I. et al. An improved smaller biotin ligase for BioID proximity labeling. Mol. Biol. Cell 27, 1188–1196 (2016).

31. May, D. G., Scott, K. L., Campos, A. R. & Roux, K. J. Comparative application of BioID and TurboID for protein-proximity biotinylation. Cells 9, 1070 (2020).

32. D’Angelo, M. A., Gomez-Cavazos, J. S., Mei, A., Lackner, D. H. & Hetzer, M. W. A change in nuclear pore complex composition regulates cell differentiation. Dev. Cell 22, 446–458 (2012).

33. Liang, Y., Franks, T. M., Marchetto, M. C., Gage, F. H. & Hetzer, M. W. Dynamic association of NUP98 with the human genome. PLOS Genet. 9, e1003308 (2013).

34. Petrovsky, R., Krohne, G. & Großhans, J. Overexpression of the lamina proteins Lamin and Kugelkern induces specific ultrastructural alterations in the morphology of the nuclear envelope of intestinal stem cells and enterocytes. Eur. J. Cell Biol. 97, 102–113 (2018).

35. Pennarun, G. et al. Increase in lamin B1 promotes telomere instability by disrupting the shelterin complex in human cells. Nucleic Acids Res. 49, 9886–9905 (2021).

36. Buchwalter, A. Intermediate, but not average: the unusual lives of the nuclear lamin proteins. Curr. Opin. Cell Biol. 84, 102220 (2023).

37. Jafferali, M. H., Figueroa, R. A., Hasan, M. & Hallberg, E. Spindle associated membrane protein 1 (Samp1) is required for the differentiation of muscle cells. Sci. Rep. 7, 16655 (2017).

38. Fu, Y. et al. MacroH2A1 associates with nuclear lamina and maintains chromatin architecture in mouse liver cells. Sci. Rep. 5, 17186 (2015).

39. Carter, S. P. et al. Genetic deletion of zebrafish Rab28 causes defective outer segment shedding, but not retinal degeneration. Front. Cell Dev. Biol. 8, 136 (2020).

40. Santoriello, C. et al. RNA helicase DDX21 mediates nucleotide stress responses in neural crest and melanoma cells. Nat. Cell Biol. 22, 372–379 (2020).

41. Sun, G. & Holley, S. A. Actn4 links inactive integrin α5 with actin in zebrafish somites. Mol. Cell. Proteomics 25, 101087 (2026).

42. Link, V., Shevchenko, A. & Heisenberg, C.-P. Proteomics of early zebrafish embryos. BMC Dev. Biol. 6, 1 (2006).

43. Purushothaman, K., et al. Proteomics analysis of early developmental stages of zebrafish embryos. Int. J. Mol. Sci. 20, 6359 (2019).

44. Karlsen, A. et al. The proteomic profile of the human myotendinous junction. iScience 25, 103836 (2022).

45. Rognoni, E., Ruppert, R. & Fässler, R. The kindlin family: functions, signaling properties and implications for human disease. J. Cell Sci. 129, 17–27 (2016).

46. Guo, L. et al. Kindlin-2 links mechano-environment to proline synthesis and tumor growth. Nat. Commun. 10, 845 (2019).

47. Liu, C. et al. Kindlin-2 enhances c-Myc translation through association with DDX3X to promote pancreatic ductal adenocarcinoma progression. Theranostics 13, 4333–4355 (2023).

48. Ma, N. et al. Kindlin-2 phase separation in response to flow controls vascular stability. Circ. Res. 135, 1141–1160 (2024).

49. Sun, Z., Costell, M. & Fässler, R. Integrin activation by talin, kindlin and mechanical forces. Nat. Cell Biol. 21, 25–31 (2019).

50. Chastney, M. R., Conway, J. R. W. & Ivaska, J. Integrin adhesion complexes. Curr. Biol. 31, R536–R542 (2021).

51. Schumacher, S., Vazquez Nunez, R., Biertümpfel, C. & Mizuno, N. Bottom-up reconstitution of focal adhesion complexes. FEBS J. 289, 3360–3373 (2022).

52. Kanchanawong, P. & Calderwood, D. A. Organization, dynamics and mechanoregulation of integrin-mediated cell–ECM adhesions. Nat. Rev. Mol. Cell Biol. 24, 142–161 (2023).

53. Li, W. et al. Kindlin2 enables EphB/ephrinB bi-directional signaling to support vascular development. Life Sci. Alliance 6, e202201800 (2023).

54. Qadota, H., Moerman, D. G. & Benian, G. M. A molecular mechanism for the requirement of PAT-4 (integrin-linked kinase (ILK)) for the localization of UNC-112 (kindlin) to integrin adhesion sites. J. Biol. Chem. 287, 28537–28551 (2012).

55. Reinhardt, S. C. M. et al. ILK binding to β1 integrin is indirectly mediated by kindlin-2. Preprint at 10.64898/2026.05.06.723202 (2026).

56. Ye, F. et al. The mechanism of kindlin-mediated activation of integrin αIIbβ3. Curr. Biol. 23, 2288–2295 (2013).

57. Li, H., et al. Structural basis of kindlin-mediated integrin recognition and activation. Proc. Natl. Acad. Sci. USA 114, 9349–9354 (2017).

58. Kadry, Y. A., Maisuria, E. M., Huet-Calderwood, C. & Calderwood, D. A. Differences in self-association between kindlin-2 and kindlin-3 are associated with differential integrin binding. J. Biol. Chem. 295, 11161–11173 (2020).

59. Jülich, D., Geisler, R. & Holley, S. A. Integrin α5 and Delta/Notch signaling have complementary spatiotemporal requirements during zebrafish somitogenesis. Dev. Cell 8, 575–586 (2005).

60. Koshida, S. et al. Integrin α5-dependent fibronectin accumulation for maintenance of somite boundaries in zebrafish embryos. Dev. Cell 8, 587–598 (2005).

61. Postel, R., Vakeel, P., Topczewski, J., Knöll, R. & Bakkers, J. Zebrafish integrin-linked kinase is required in skeletal muscles for strengthening the integrin–ECM adhesion complex. Dev. Biol. 318, 92–101 (2008).

62. Dowling, J. J. et al. Kindlin-2 is an essential component of intercalated discs and is required for vertebrate cardiac structure and function. Circ. Res. 102, 423–431 (2008).

63. Kirchmaier, B. C. et al. The Popeye domain containing 2 (popdc2) gene in zebrafish is required for heart and skeletal muscle development. Dev. Biol. 363, 438–450 (2012).

64. Jacob, A. E., Amack, J. D. & Turner, C. E. Paxillin genes and actomyosin contractility regulate myotome morphogenesis in zebrafish. Dev. Biol. 425, 70–84 (2017).

65. Han, M. K. L., Van Der Krogt, G. N. M. & De Rooij, J. Zygotic vinculin is not essential for embryonic development in zebrafish. PLOS ONE 12, e0182278 (2017).

66. Hunter, M. J. & Ludwig, M. L. The reaction of imidoesters with proteins and related small molecules. J. Am. Chem. Soc. 84, 3491–3504 (1962).

67. Smith, A. L., Friedman, D. B., Yu, H., Carnahan, R. H. & Reynolds, A. B. ReCLIP (reversible cross-link immuno-precipitation): an efficient method for interrogation of labile protein complexes. PLOS ONE 6, e16206 (2011).

68. Huang, H.-Y. et al. Tdrd1 acts as a molecular scaffold for Piwi proteins and piRNA targets in zebrafish. EMBO J. 30, 3298–3308 (2011).

69. Roovers, E. F. et al. Tdrd6a regulates the aggregation of Buc into functional subcellular compartments that drive germ cell specification. Dev. Cell 46, 285–301.e9 (2018).

70. Sciuto, M. R. et al. Two-step coimmunoprecipitation (TIP) enables efficient and highly selective isolation of native protein complexes. Mol. Cell. Proteomics 17, 993–1009 (2018).

71. Papachristou, E. K. et al. A quantitative mass spectrometry-based approach to monitor the dynamics of endogenous chromatin-associated protein complexes. Nat. Commun. 9, 2311 (2018).

72. Levic, D. S., Yamaguchi, N., Wang, S., Knaut, H. & Bagnat, M. Knock-in tagging in zebrafish facilitated by insertion into non-coding regions. Development 148, dev199994 (2021).

73. Shen, J. et al. Zebrafish knock-in lines enabling live visualization of extracellular matrix dynamics during development and regeneration. Development 153, dev205402 (2026).

74. Simsek, M. F. et al. The vertebrate segmentation clock drives segmentation by stabilizing Dusp phosphatases in zebrafish. Dev. Cell 60, 669–678.e6 (2025).

