## Supplementary Figures 1-13 for "SureCLIP and DrCLIP: Genetic manipulation-free, sub-hour protein interaction mapping in cells and zebrafish"

### Supplementary Figure 1

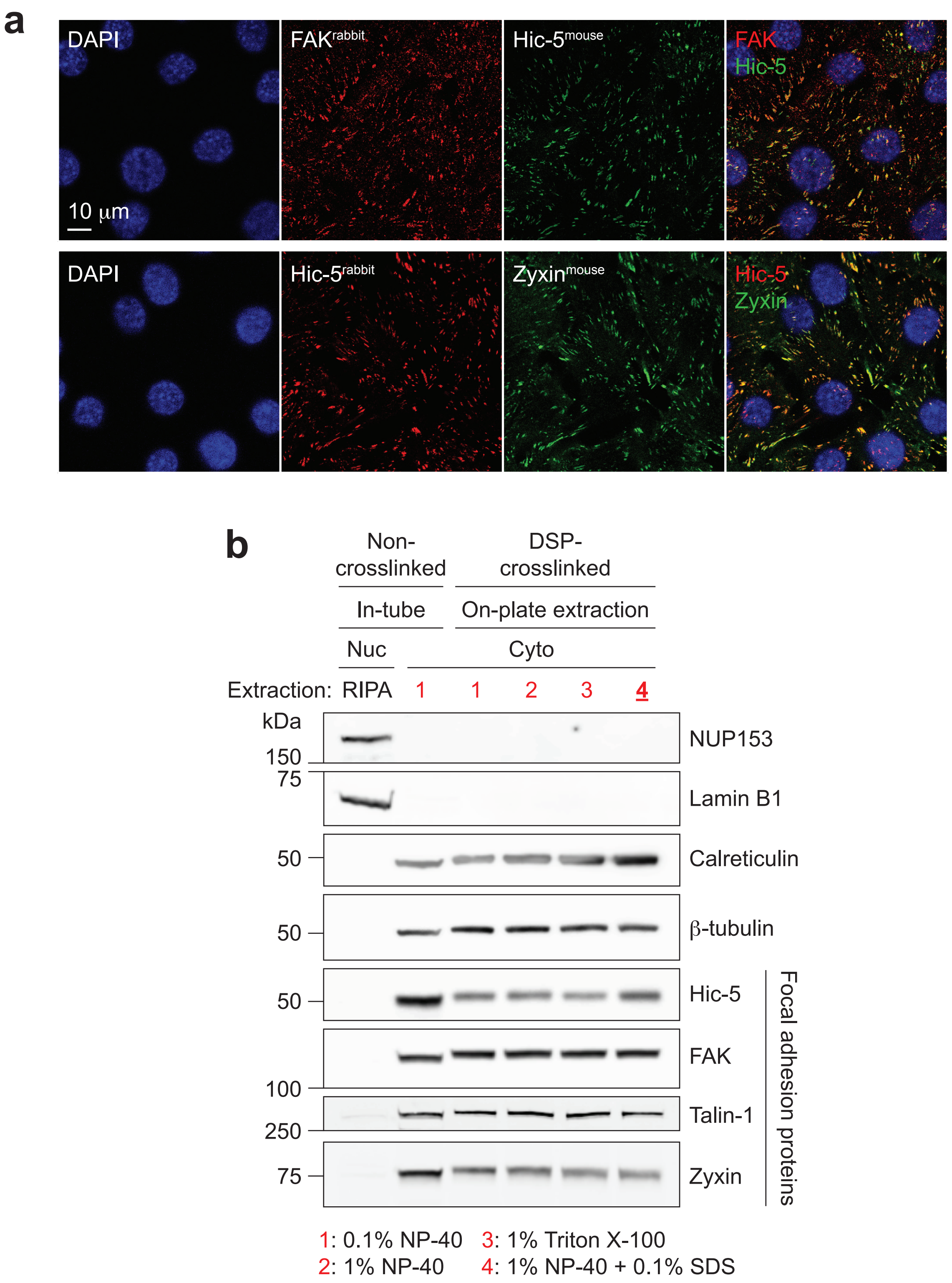


**Supplementary Figure 1. Subcellular localization and fractionation profile of Hic-5.** (**a**) Immunostaining of Hic-5, FAK, and zyxin in C2C12 cells. Superscript indicates the antibody host. (**b**) Cytoplasmic extraction from non- and DSP-crosslinked C2C12 cells. Nuc: nuclear; Cyto: cytoplasmic. All subsequent SureCLIP experiments used condition 4.

### Supplementary Figure 2

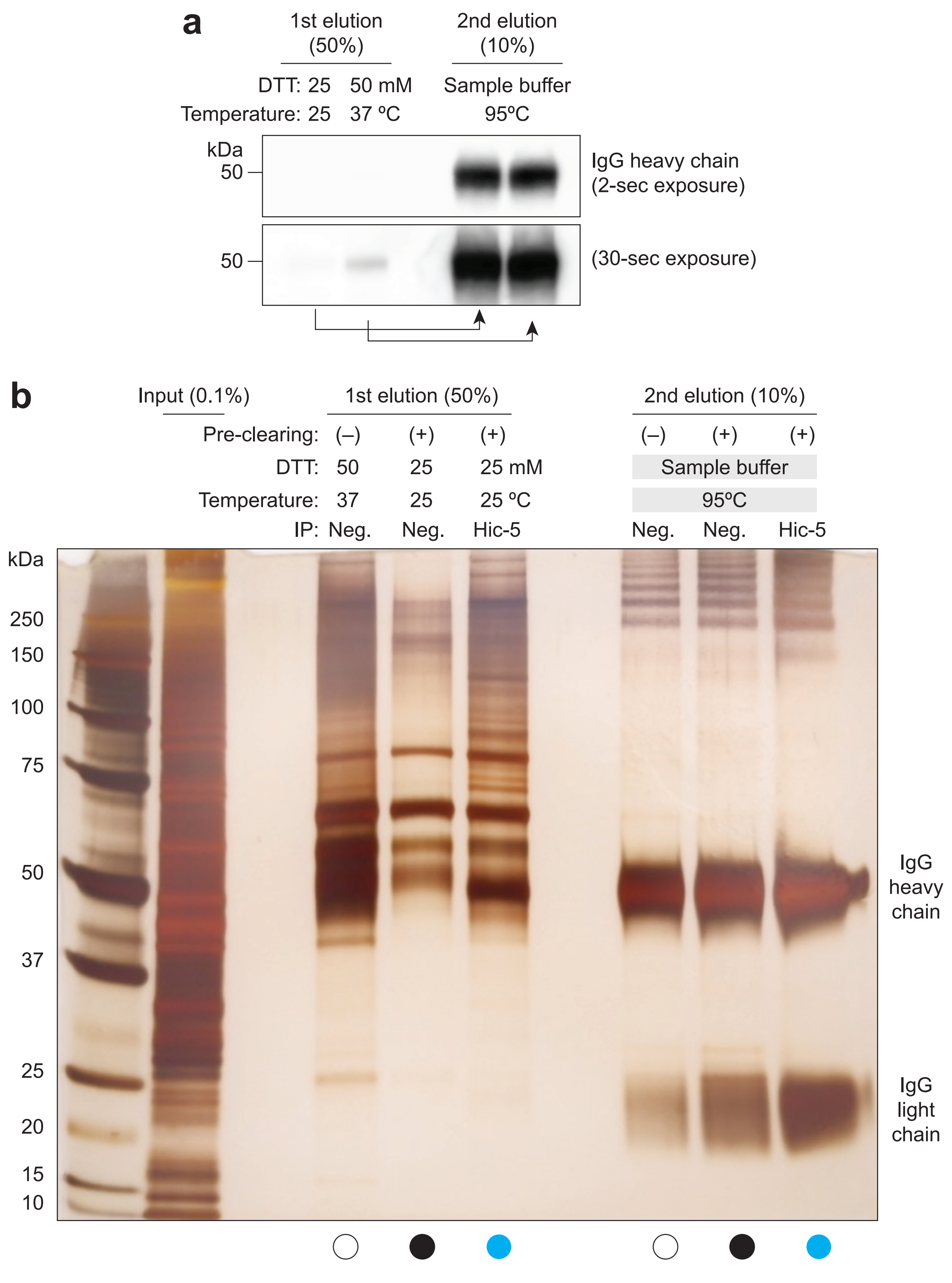


**Supplementary Figure 2. SureCLIP elution protocol minimizes IgG contamination.** (**a**) Dissociation of IgG heavy chain from Protein A beads was assessed under two different temperatures and DTT concentrations. After incubating in DTT-supplemented RIPA buffer for 30 min (1st elution), beads were added 5x sample buffer and heated at 95°C for 4 min (2nd elution). (**b**) Samples from Fig. 1d were run under identical conditions on an SDS-PAGE gel and silver stained.

### Supplementary Figure 3

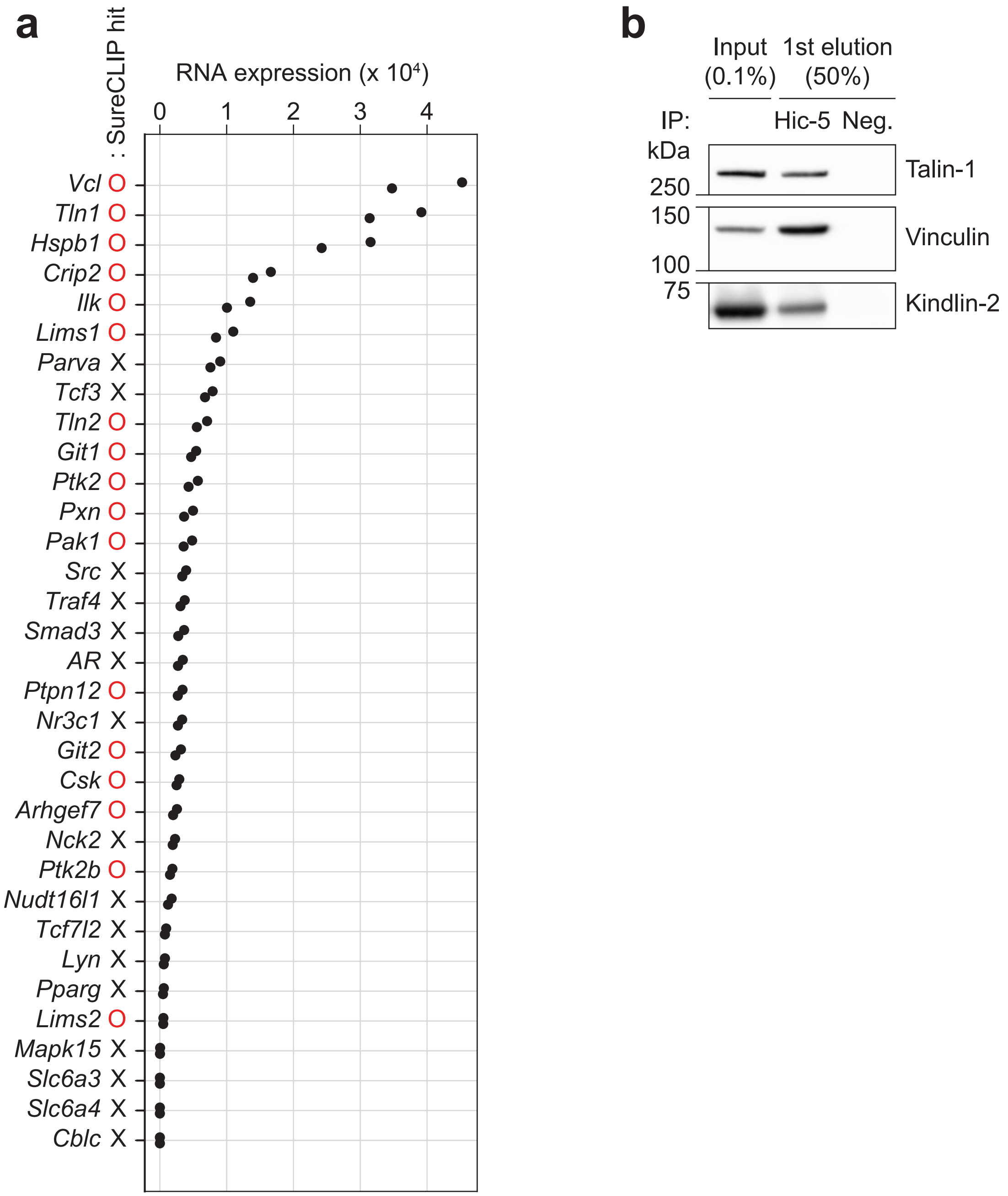


**Supplementary Figure 3. Hic-5 interactors identified by SureCLIP.** (**a**) RNA expression levels of UniProt-annotated Hic-5 interactors in C2C12 myoblasts. RNA-sequencing data source: GEO database GSE183521. (**b**) Immunoblot analysis of the input and first eluate from Hic-5 SureCLIP, probed for talin-1, vinculin, and kindlin-2.

### Supplementary Figure 4

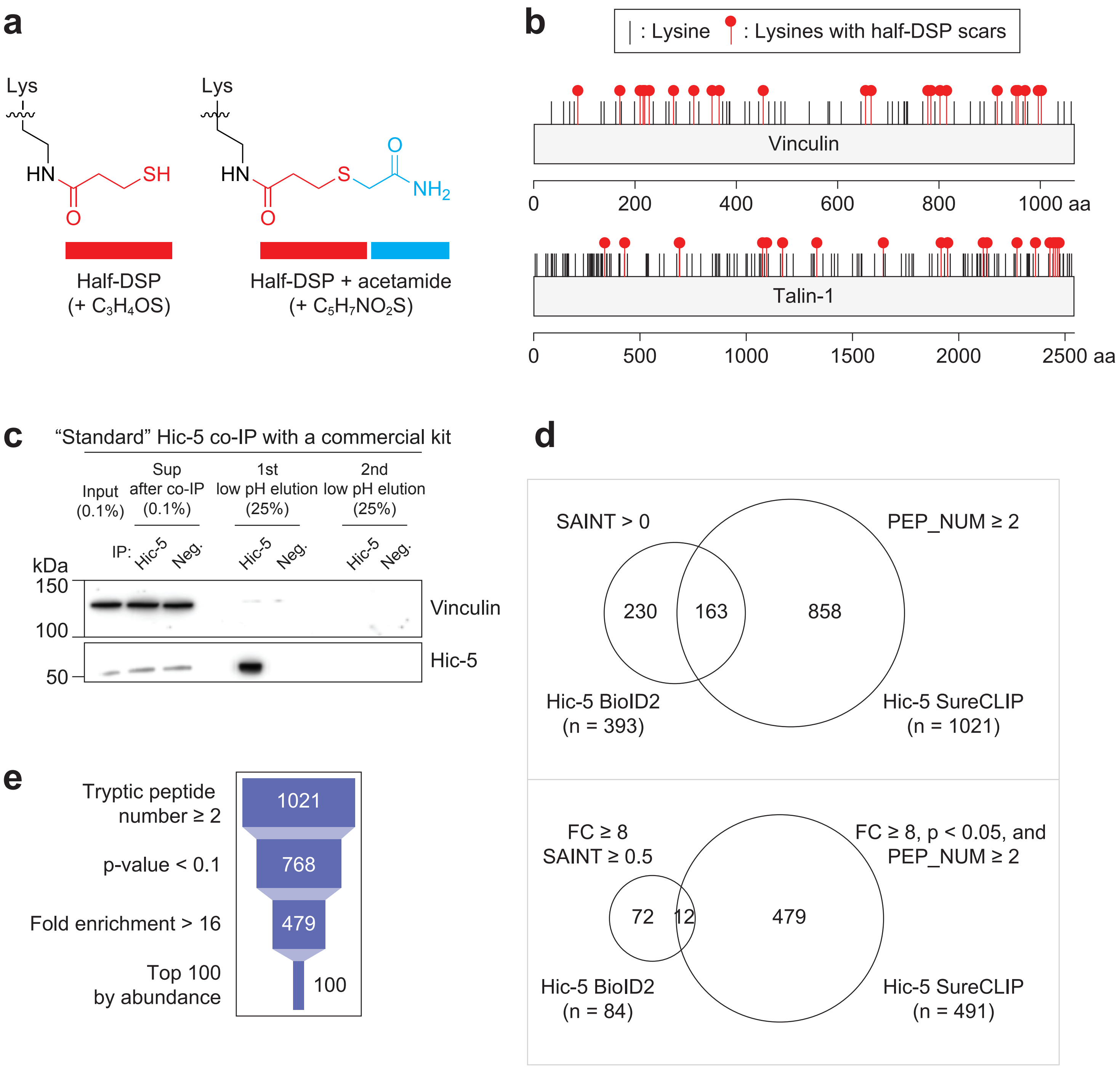


**Supplementary Figure 4. SureCLIP leaves defined DSP scars and outperforms standard co-IP.** (**a**) Chemical structures of lysine residues modified with half-DSP adducts. Acetamide moiety is derived from chloro/iodoacetamide added during the mass spectrometry sample preparation step. (**b**) Lysine residues in vinculin and talin-1 that reacted with DSP. aa: amino acid. (**c**) "Standard" Hic-5 co-IP was carried out using a commercial kit. The levels of vinculin and Hic-5 in the input, post-co-IP supernatant, and two sequential eluates were analyzed by immunoblotting. Proteins were eluted from the beads using a pH 2.8 buffer. (**d**) Overlap of Hic-5 interactors identified by BioID2 and SureCLIP. SAINT: Significance Analysis of INTeractome. (**e**) Criteria used to filter SureCLIP hits.

### Supplementary Figure 5

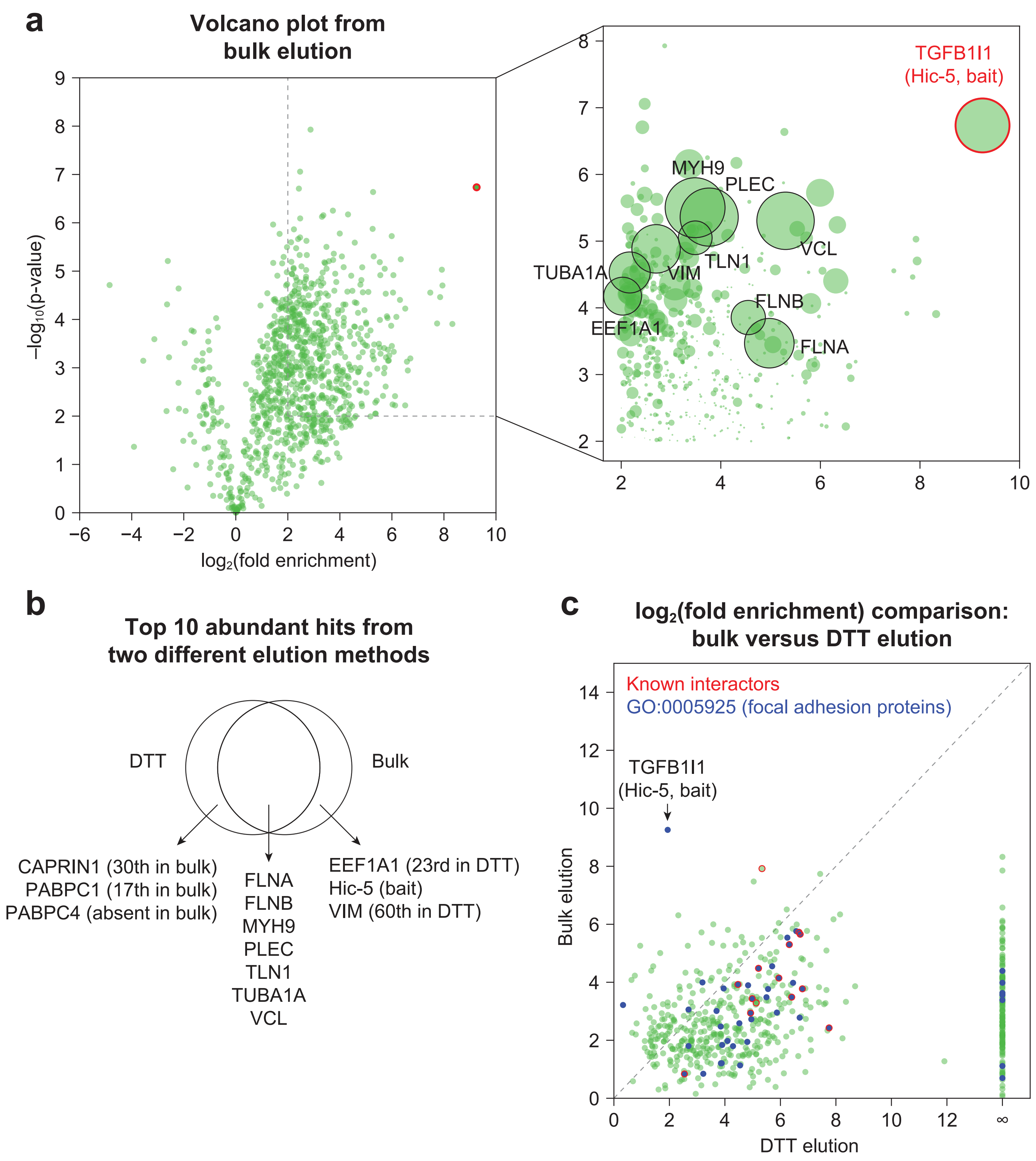


**Supplementary Figure 5. Comparison of bulk and DTT-based elution in SureCLIP.** (**a**) Volcano plot from Hic-5 SureCLIP with bulk elution. In the zoomed view, datapoint size reflects protein abundance. (**b**) Overlap of the 10 most abundant hits (> 4-fold enrichment, $p < 0.05$) from DTT and bulk elution. Abundance ranks in the other data-set are in parentheses. (**c**) Fold enrichment comparison of bulk versus DTT elution. Red, known Hic-5 interactors; blue, focal adhesion proteins (GO:0005925). Dashed line, $y = x$.

### Supplementary Figure 6

**a**

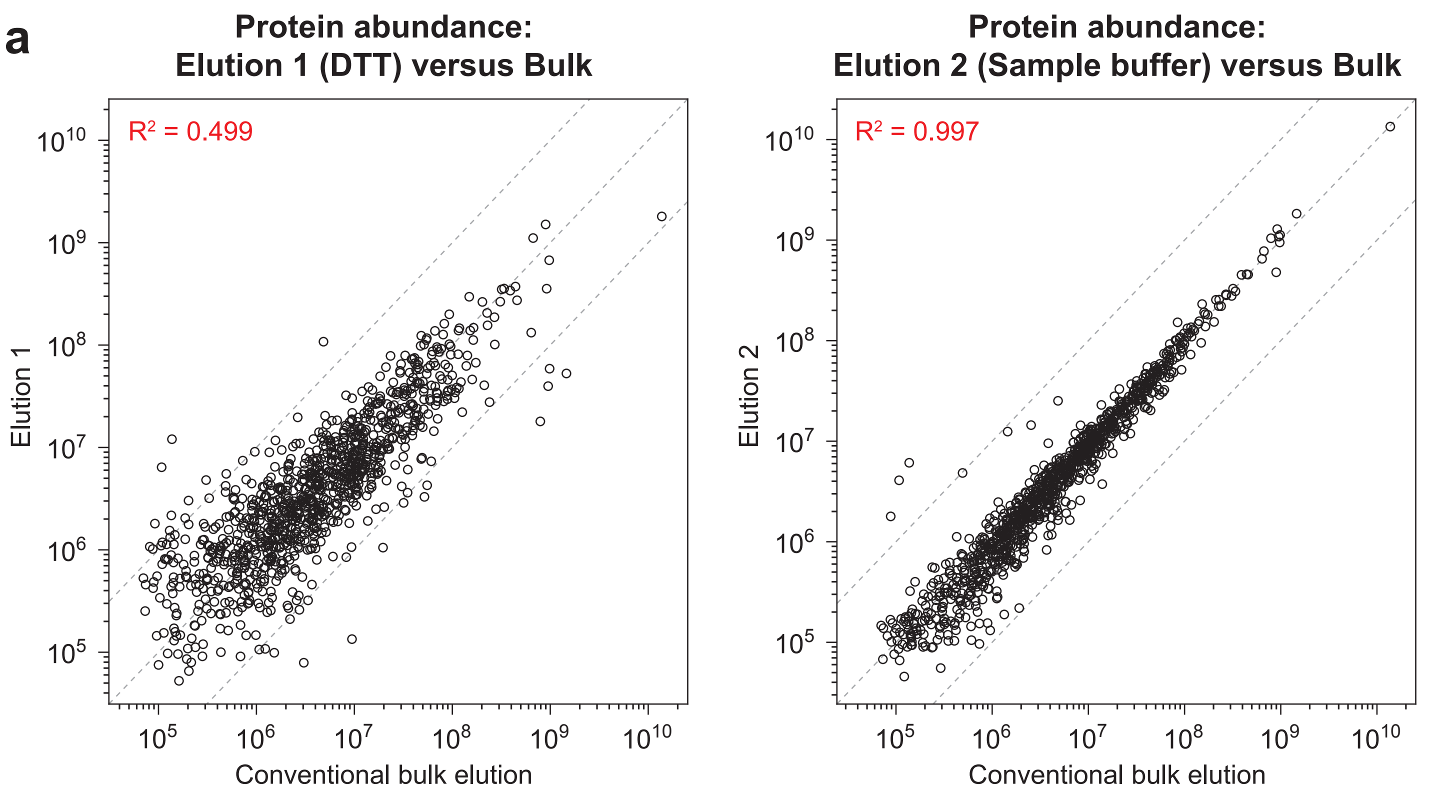


**b**

| | log₂(fold enrichment versus negative control) |
|---|---|
| NEFL | 4.7 |
| TGFB1I1 | 9.3 |
| SNRPF | -1.6 |
| SNRPD1 | -0.1 |
| NUCB1 | 1.2 |
| AKAP9 | 2.2 |
| SNRNP70 | 1.0 |
| NPEPPS | 0.8 |
| ITPR3 | 3.5 |
| RPL30 | 0.9 |
| KRT1 | 0.1 |
| PRPF38A | 0.7 |
| GNAS | 2.4 |
| KPNA3 | -0.9 |
| EHD2 | 1.7 |
| RALY | 3.3 |
| SNRPE | -2.2 |
| SNRPD3 | -1.4 |
| AP3B1 | 3.9 |
| DES | 3.4 |
| H4C1 | -0.2 |
| CFAP20 | -0.9 |
| CLPX | -0.3 |
| RPL22L1 | 1.3 |
| ATP5PO | 1.0 |
| MYOF | 0.4 |
| NEFM | 6.5 |
| KRT81 | 1.0 |
| RTN4 | 0.7 |
| AGAP3 | 0.3 |

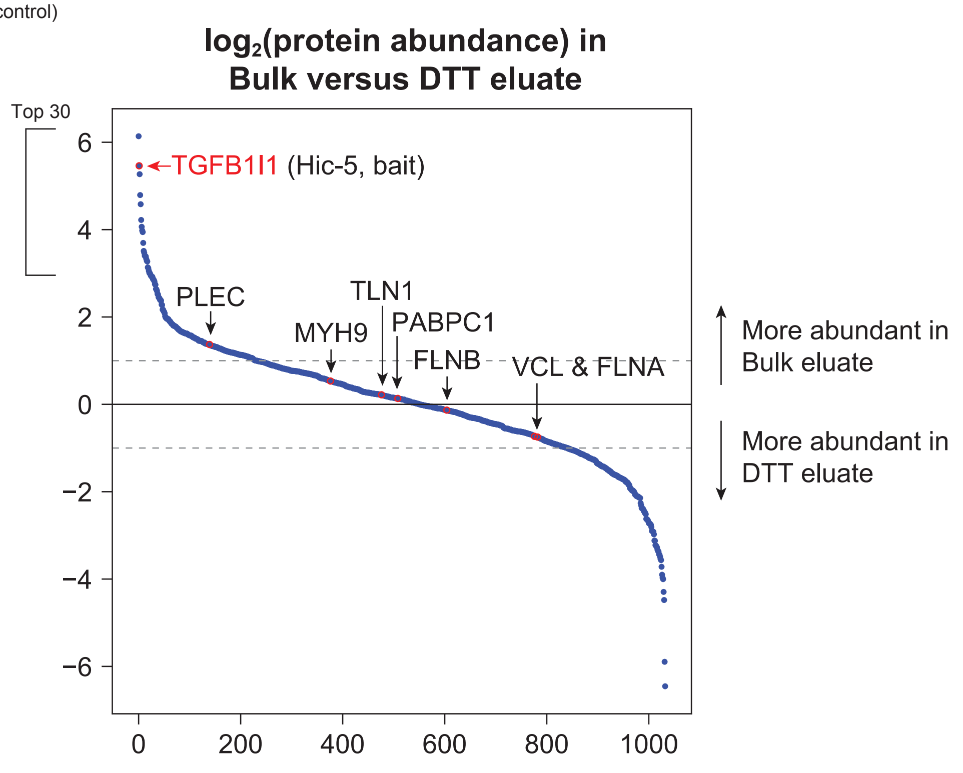


**Supplementary Figure 6. Bulk elution recovers nonspecific material excluded by DTT elution.** (**a**) Protein abundance in the first (DTT, left) and second (sample buffer, right) eluates versus the bulk eluate. Dashed lines, y = x and 10-fold deviation. (**b**) Proteins ranked by $log_2$(abundance ratio) of bulk versus DTT eluate. Dashed lines, 2-fold difference. Left, $log_2$(fold enrichment over control) in the bulk eluate for the top 30 proteins more abundant in the bulk eluate.

### Supplementary Figure 7

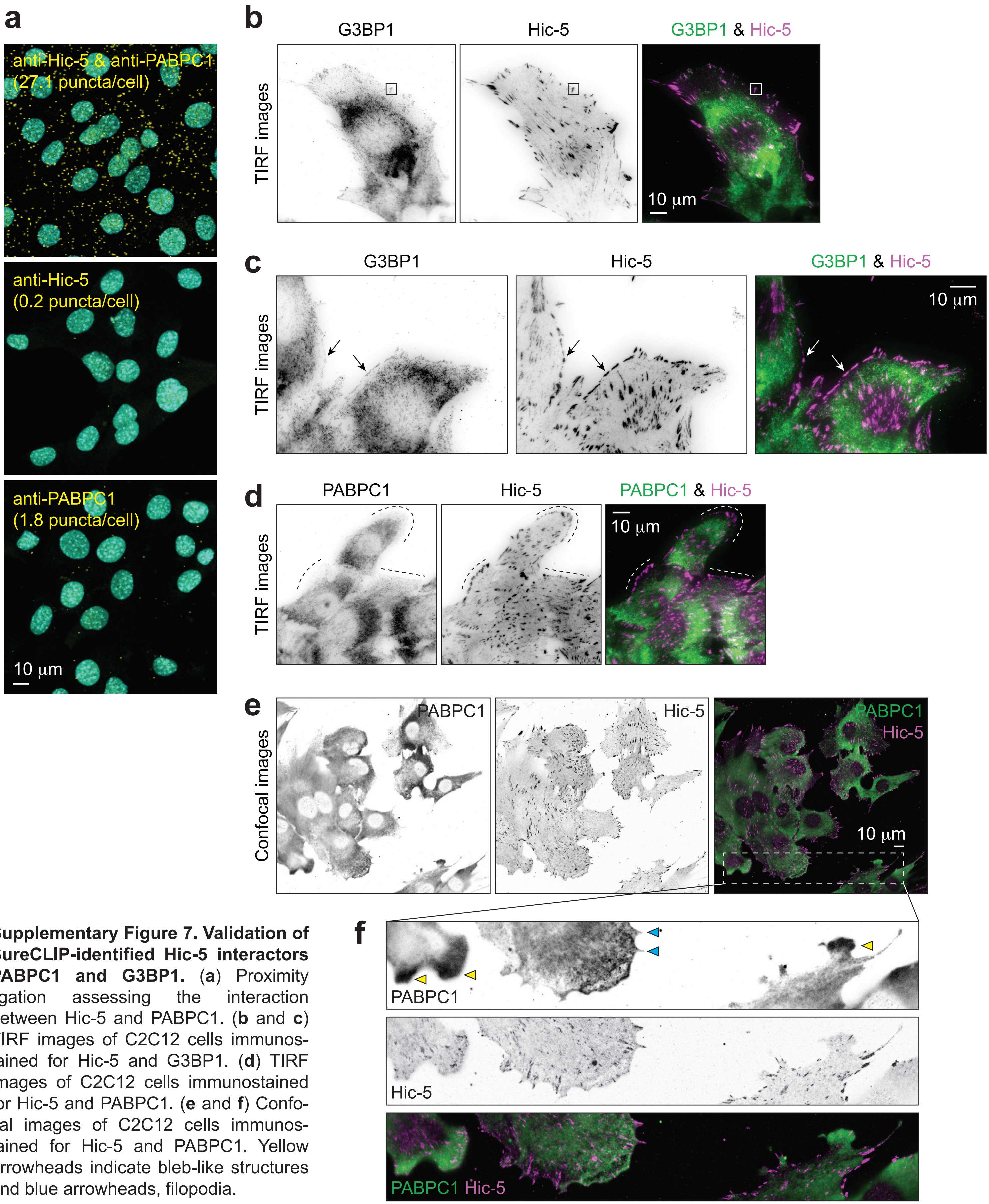


**Supplementary Figure 7. Validation of SureCLIP-identified Hic-5 interactors PABPC1 and G3BP1.** (**a**) Proximity ligation assessing the interaction between Hic-5 and PABPC1. (**b** and **c**) TIRF images of C2C12 cells immunostained for Hic-5 and G3BP1. (**d**) TIRF images of C2C12 cells immunostained for Hic-5 and PABPC1. (**e** and **f**) Confocal images of C2C12 cells immunostained for Hic-5 and PABPC1. Yellow arrowheads indicate bleb-like structures and blue arrowheads, filopodia.

### Supplementary Figure 8

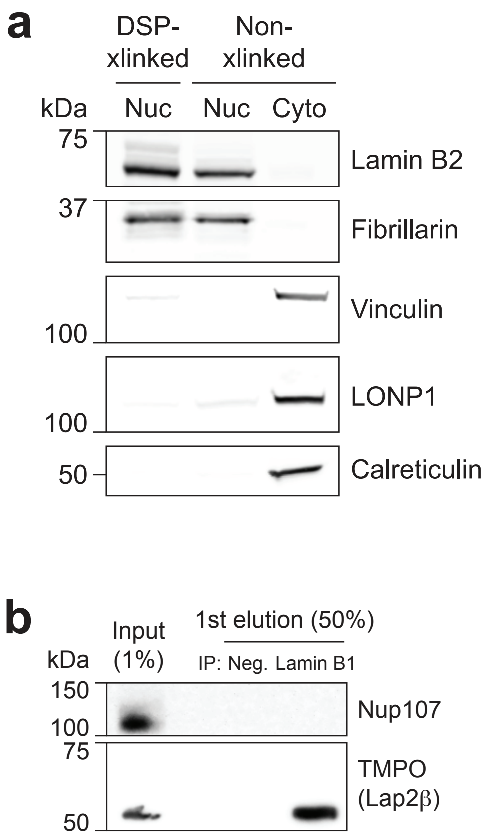


**Supplementary Figure 8. SureCLIP with lamin B1 as bait.** (**a**) Nuclear extracts prepared from C2C12 cells with or without DSP crosslinking. (**b**) Immunoblot validation of LAP2 in the first (DTT) eluate from lamin B1 SureCLIP. Nup107 serves as a negative control. Neg.: negative (IgG isotype control).

### Supplementary Figure 9

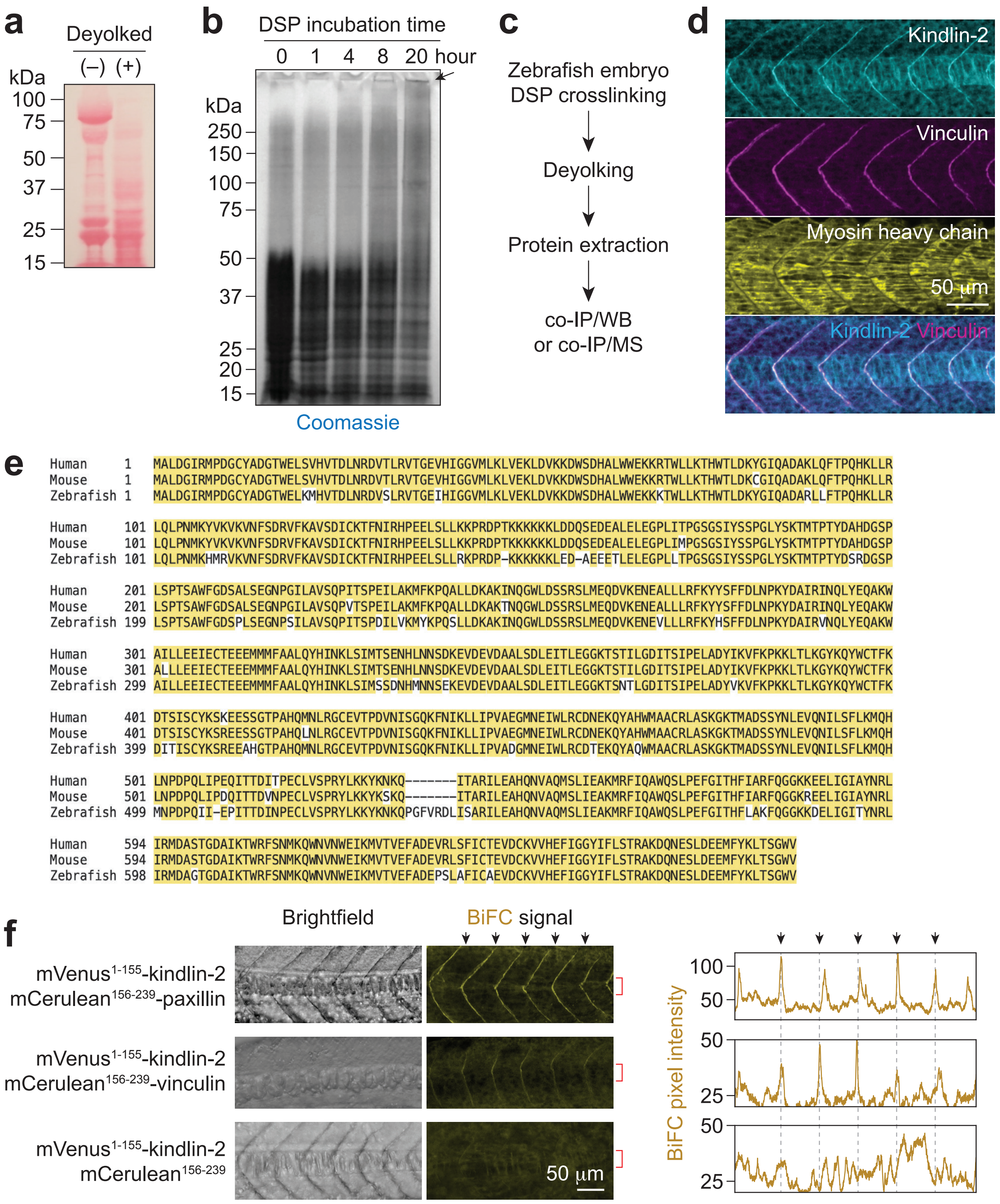


**Supplementary Figure 9. Optimization of DrCLIP and validation of kindlin-2 interactions in zebrafish embryos.** (**a**) Ponceau S staining of lysates from deyolked and non-deyolked 24-hpf embryos. (**b**) Embryos at 24 hpf were DSP-crosslinked for 0, 1, 4, 8, or 20 hours and then deyolked. Resulting lysates were analyzed by non-reducing SDS-PAGE and Coomassie staining. (**c**) Schematic of the DrCLIP workflow. (**d**) Immunostaining of kindlin-2, vinculin, and myosin heavy chain in 24-hpf embryo somites. (**e**) Alignment of human, mouse, and zebrafish kindlin-2 protein sequences. (**f**) BiFC analysis of kindlin-2/paxillin and kindlin-2/vinculin interactions in 24-hpf embryos. BiFC intensity profiles along the somite anterior–posterior axis were measured in the regions marked by red brackets. Arrows indicate corresponding somite positions.

### Supplementary Figure 10

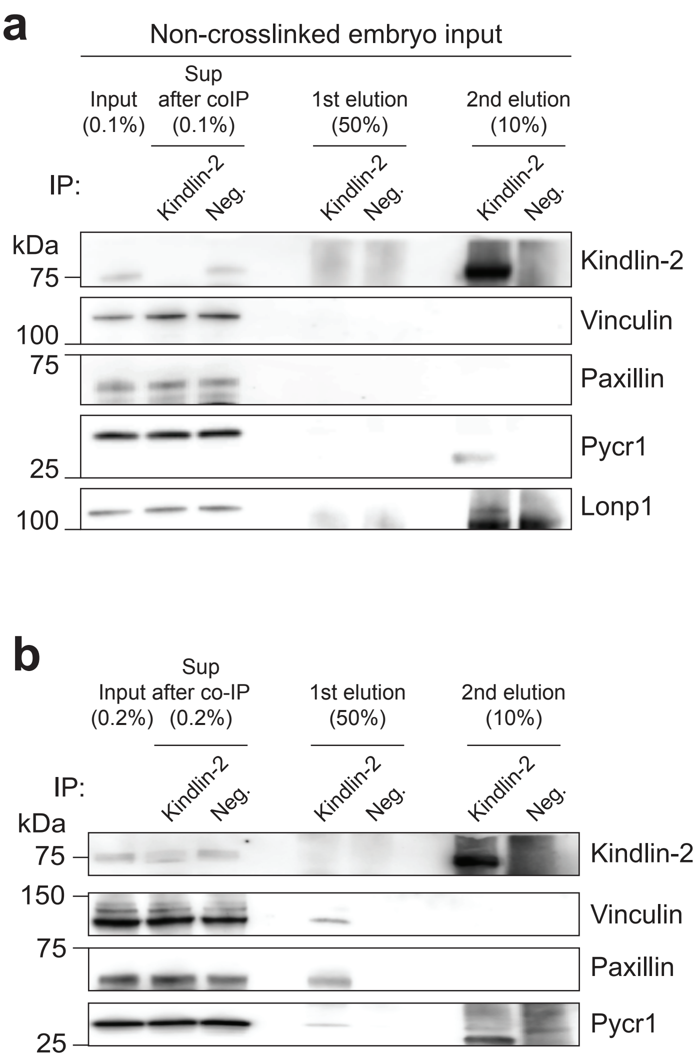


**Supplementary Figure 10. Crosslinker dependence of DrCLIP.** (**a**) DrCLIP was performed on lysates from non-crosslinked 24-hpf embryos. (**b**) DrCLIP was performed on 3-dpf larvae crosslinked with 5 mM DTBP for 1 hour. In panels **a** and **b**, kindlin-2 and its interactors were detected by immunoblotting in the input, post-co-IP supernatant, and eluates.

### Supplementary Figure 11

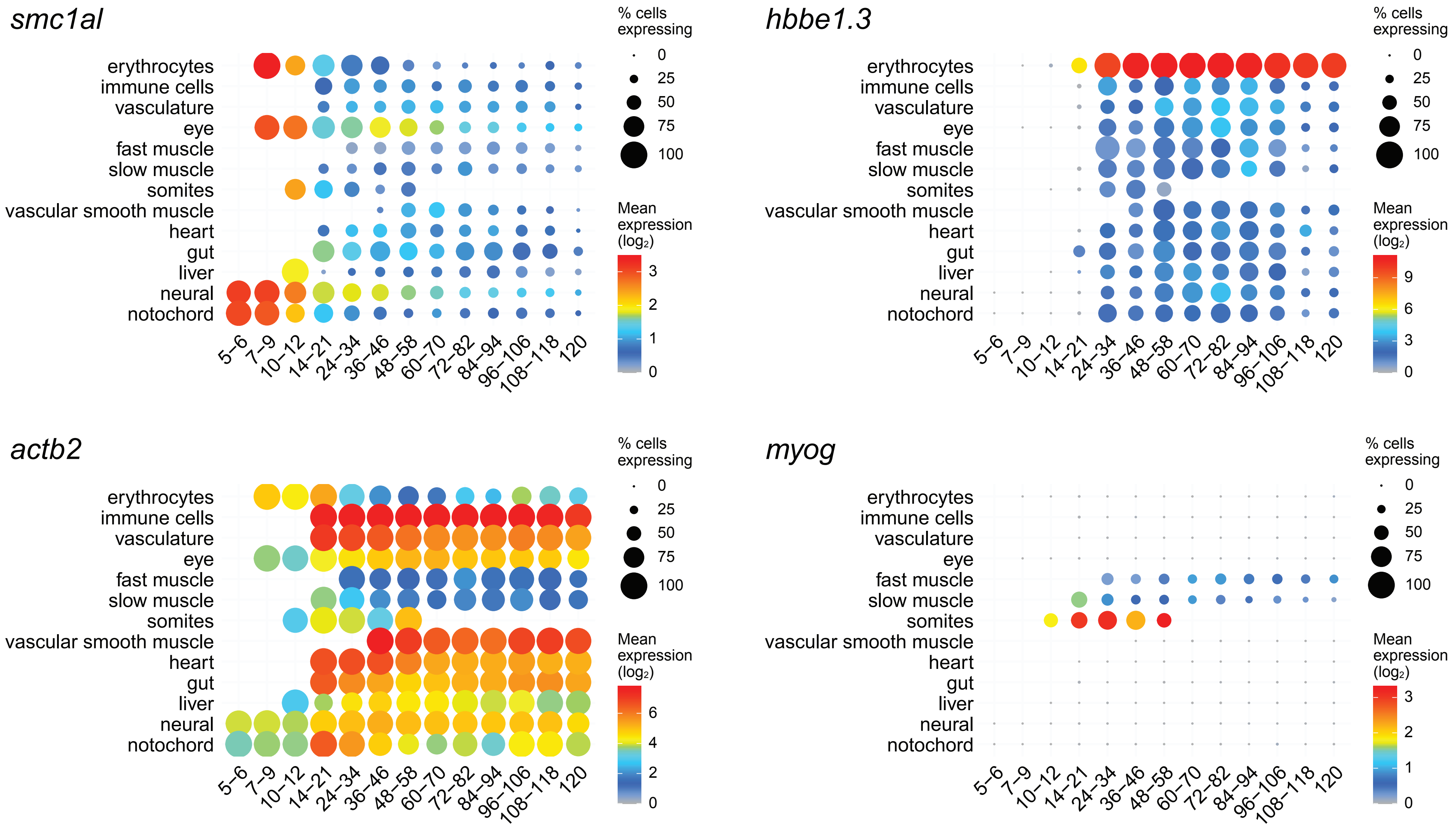


**Supplementary Figure 11. Developmental expression of selected genes from the Daniocell atlas.** *actb2* and *myog* are shown as examples of ubiquitously expressed and tissue-specific genes, respectively. Note that color scales differ among the four genes. x-axis: hpf. (Daniocell version 1.1; https://daniocell.nichd.nih.gov)

### Supplementary Figure 12

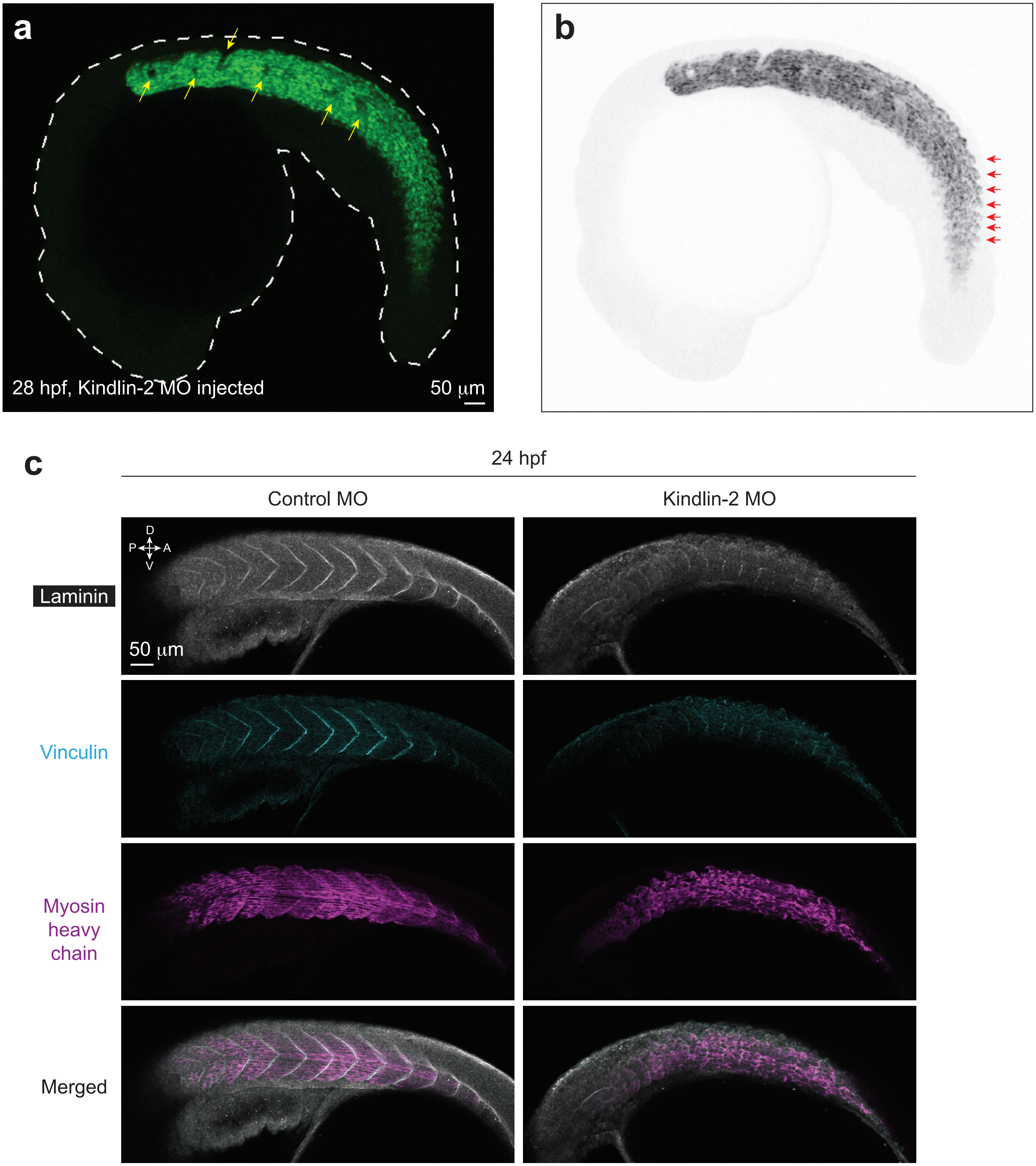


**Supplementary Figure 12. Kindlin-2 is required for ECM assembly at somite boundaries but not for segmentation.** (**a**) Immunostaining of myosin heavy chain in a kindlin-2 morphant embryo at 28 hpf. The dashed line outlines the embryo. Yellow arrows mark gaps in the trunk musculature. (**b**) Inverted grayscale image of (**a**). Red arrows mark segmented somites in the posterior trunk. (**c**) Immunostaining of laminin, vinculin, and myosin heavy chain in control and kindlin-2 morphant embryos at 24 hpf. A, anterior; P, posterior; D, dorsal; V, ventral.

### Supplementary Figure 13

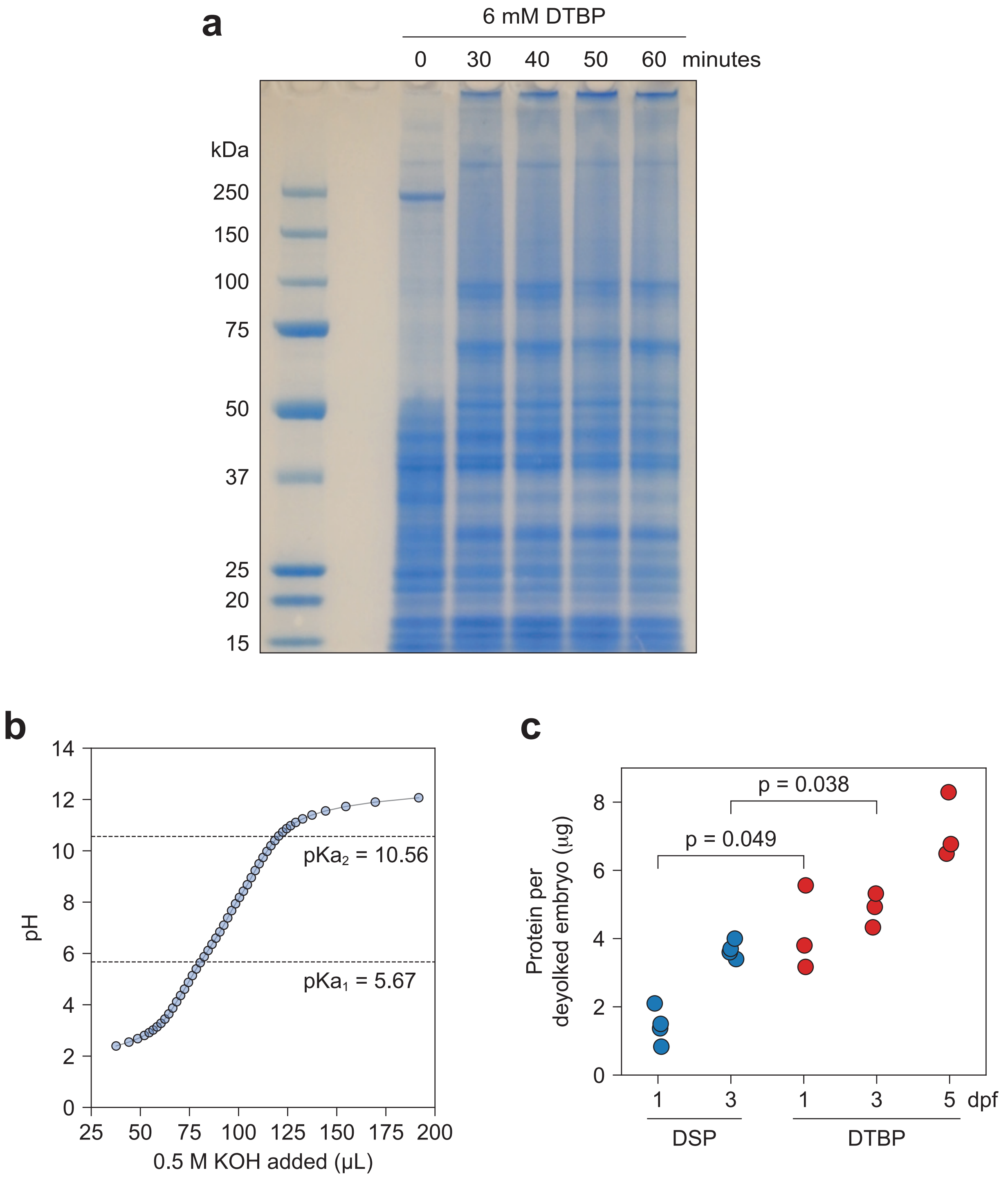


**Supplementary Figure 13. DTBP crosslinking kinetics and protein extraction from crosslinked embryos.** (**a**) 3-dpf larvae were crosslinked with 6 mM DTBP for the indicated durations, lysed, and resolved by non-reducing SDS-PAGE. (**b**) Titration of DTBP·2HCl with KOH. (**c**) Protein yield per deyolked embryo by time point and crosslinker. Each dot, mean of at least 30 embryos.
