## Supplementary Table 7 for "SureCLIP and DrCLIP: Genetic manipulation-free, sub-hour protein interaction mapping in cells and zebrafish"

**ANTIBODIES**

| **Target (host)** | **Manufacturer** | **Catalog #** | **Application*** |
| --- | --- | --- | --- |
| Hic-5 (rabbit) | Proteintech | 10565-1-AP | WB, IF, IP, and PLA |
| Hic-5 (mouse) | BD Transduction | 611164 | WB and IF |
| Kindlin-2 | Proteintech | 11453-1-AP | WB, IF, and IP |
| Lamin B1 | Proteintech | 66095-1-Ig | WB and IP |
| Lamin B2 | Proteintech | 10895-1-AP | WB |
| Histone H2B | Proteintech | 68393-1-Ig | WB |
| Fibrillarin | Cell Signaling Technology | 2639S | WB |
| Vinculin | Proteintech | 66305-1-Ig | WB |
| Vinculin | Sigma | V9131 | WB and IF |
| Laminin | Sigma | L9393 | IF |
| Paxillin | BD Biosciences | 610051 | WB and IF |
| Talin-1 | Cell Signaling Technology | 4021S | WB |
| Zyxin | R&D Systems | MAB6977 | WB and IF |
| FAK (rabbit) | Cell Signaling Technology | 3285T | WB and IF |
| FAK (mouse) | BD Transduction | 610087 | WB |
| PYCR1 | Proteintech | 13108-1-AP | WB |
| β-tubulin | Proteintech | 66240-1-Ig | WB |
| LONP1 | Cell Signaling Technology | 28020S | WB |
| Calreticulin | Cell Signaling Technology | 12238S | WB |
| EEF2 | Proteintech | 67550-1-Ig | WB |
| PABPC1 | Proteintech | 66809-1-Ig | WB, IF, and PLA |
| G3BP1 | Proteintech | 66486-1-Ig | WB, IF, and PLA |
| Caspase-3 | Cell Signaling Technology | 9662S | WB |
| Rabbit IgG Control | Proteintech | 30000-0-AP | IP |
| Mouse IgG Control | Proteintech | B900620 | IP |

* WB: western blotting, IF: immunofluorescence, IP: immunoprecipitation, and PLA: proximity ligation assay

**REAGENTS**

| **Reagent** | **Manufacturer** | **Cat #** |
| --- | --- | --- |
| Protein A beads | Invitrogen | 10002D |
| Protein G beads | Invitrogen | 10003D |
| Silver stain kit | Thermo Scientific | 24612 |
| Coomassie | Bio-Rad | 1610786 |
| Ponceau S | Thermo Scientific | A40000279 |
| Pre-cast Bis-Tris gels | Invitrogen | NW04122BOX |
| SP6 in vitro transcription kit | Invitrogen | AM1340 |
| DSP | ProteoChem | c1106 |
| DTBP | Thermo Scientific | 20665 |
| Bicine buffer (pH 8.5) | Thermo | J61632.AE |
| Formaldehyde | Thermo Scientific | 28908 |
| Bovine serum albumin | Prometheus | 25-529 |
| Pierce Crosslink Magnetic IP/co-IP Kit | Thermo Scientific | 88805 |
