## Supplementary figures and images for "SureCLIP and DrCLIP: Genetic manipulation-free, sub-hour protein interaction mapping in cells and zebrafish"

### Supplementary Video 2

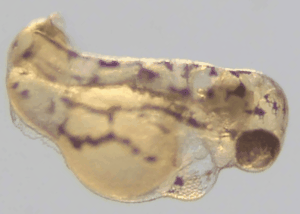
